# The genome of *Peronospora belbahrii* reveals high heterozygosity, a low number of canonical effectors and CT-rich promoters

**DOI:** 10.1101/721027

**Authors:** Marco Thines, Rahul Sharma, Sander Y. A. Rodenburg, Anna Gogleva, Howard S. Judelson, Xiaojuan Xia, Johan van den Hoogen, Miloslav Kitner, Joël Klein, Manon Neilen, Dick de Ridder, Michael F. Seidl, Guido Van den Ackerveken, Francine Govers, Sebastian Schornack, David J. Studholme

## Abstract

Along with *Plasmopara destructor, Peronosopora belbahrii* has arguably been the economically most important newly emerging downy mildew pathogen of the past two decades. Originating from Africa, it has started devastating basil production throughout the world, most likely due to the distribution of infested seed material. Here we present the genome of this pathogen and results from comparisons of its genomic features to other oomycetes. The assembly of the nuclear genome was ca. 35.4 Mbp in length, with an N50 scaffold length of ca. 248 kbp and an L50 scaffold count of 46. The circular mitochondrial genome consisted of ca. 40.1 kbp. From the repeat-masked genome 9049 protein-coding genes were predicted, out of which 335 were predicted to have extracellular functions, representing the smallest secretome so far found in peronosporalean oomycetes. About 16 % of the genome consists of repetitive sequences, and based on simple sequence repeat regions, we provide a set of microsatellites that could be used for population genetic studies of *Pe. belbahrii. Peronospora belbahrii* has undergone a high degree of convergent evolution, reflecting its obligate biotrophic lifestyle. Features of its secretome, signalling networks, and promoters are presented, and some patterns are hypothesised to reflect the high degree of host specificity in *Peronospora* species. In addition, we suggest the presence of additional virulence factors apart from classical effector classes that are promising candidates for future functional studies.

## Introduction

Oomycetes comprise more than 2000 species with a broad array of lifestyles. They are found in almost all ecosystems, efficiently colonising decaying organic matter as well as living hosts and play important ecological roles as saprotrophs and pathogens (Judelson 2012; Thines 2014). Many oomycete species are responsible for huge economic losses and pose a major threat to food security. Unlike fungi, oomycetes are diploid for most of their life cycle and therefore have the potential to propagate as heterozygous clones.

The majority of known oomycete species causes downy mildew and is obligate biotrophic. Among this group of more than 800 species, about 500 belong to the genus *Peronospora* (Thines 2014; Thines and Choi 2016). *Peronospora belbahrii* is a highly specialised species, causing basil downy mildew disease (Thines et al. 2009), a major threat to production of the culinary herb basil *(Ocimum basilicum).* The lifestyle of pathogens is reflected by its primary and secondary metabolism. Obligate biotrophs fully rely on their host for survival and utilize metabolites provided by their hosts (Judelson 2017). Therefore, obligate biotrophs have reduced sets of enzyme-encoding genes required for metabolic pathways (Baxter et al. 2010; Kemen et al. 2011; Sharma et al. 2015a, b; Spanu 2012). For example, in addition to the widespread auxotrophy for thiamine and sterol in oomycetes (Dahlin et al. 2017; Judelson 2017), obligate biotrophic oomycetes have lost the ability to reduce inorganic nitrogen and sulphur (Baxter et al. 2010; Kemen et al. 2011; Sharma et al. 2015a, b; Spanu 2012).

The first line of pathogen attack relies on hydrolytic enzymes such as cutinases, glycosyl hydrolases, lipases, and proteases that degrade plant cell wall components to promote pathogen entry (Blackman et al. 2015). Besides these enzymes, oomycetes secrete effectors that target host processes (Schornack et al. 2009). After germination and penetration, downy mildews are nutritionally dependent on living host cells from which they extract nutrients *via* haustoria, which are believed to represent the main interaction interface between host and pathogen (Schornack et al. 2009; Ellis et al. 2007; Soylu and Soylu 2003). Extracellular proteins secreted form hyphae and haustoria play key roles in establishing stable interactions by modulating plant immune responses and plant metabolism (Meijer et al. 2014). While the biological function of many oomycete effectors still remains to be resolved, they have been classified based on sequence similarity and other recognizable sequence features such as the WY domain (Boutemy et al. 2011) into host-translocated (cytoplasmic) effectors and apoplastic effectors. The latter include protease inhibitors, small cysteine rich proteins (SCPs), elicitins and necrosis-inducing-like protein (NLPs). NLPs are widely distributed across plant pathogenic oomycetes and share a highly conserved NPP1 domain (Feng et al. 2014; Seidl and Ackerveken 2019). Most cytoplasmic effectors carry short motifs associated with their translocation, such as the characteristic RxLR effector motif that often cooccurs with a downstream EER motif (Dou et al. 2008; Tyler et al. 2013).

The availability of next-generation sequencing enables economical and facile analysis of genomes, leading to many important insights into the biology of many oomycetes (Baxter et al. 2010; Ali et al. 2017; McCarthy and Fitzpatrick 2017; Lamour et al. 2012; Raffaele and Kamoun 2012; Tyler et al. 2006; Lévesque et al. 2010; Tian et al. 2011; Soanes et al. 2007; Sharma et al. 2015; Judelson 2012; Grünwald 2012; Jiang et al. 2013; Links et al. 2011; Haas et al. 2009; Schena et al. 2008; Gaulin et al. 2018). Because of their economic importance, some downy mildews have had their genomes sequenced, including: *Bremia lactucae, Hyaloperonospora arabidopsidis, Peronospora effusa, Peronospora tabacina, Plasmopara viticola, Plasmopara halstedii, Plasmopara muralis, Plasmopara obducβns, Pseuoperonospora cubensis, Pseudoperonospora humuli,* and *Sclerospora graminicola* (Fletcher et al. 2019; Baxter et al. 2010; Feng et al. 2018; Fletcher et al. 2018; Derevnina et al. 2015; Sharma et al. 2015; Kobayashi et al. 2017; Tian et al. 2011; Ye et al. 2016; Nayaka et al. 2017, respectively). Here we augment this repertoire with the annotated genome sequence of *Pe. belbahrii.* Using comparative genomics we uncover evidence of convergent evolution in loss of metabolic function between this species and phylogenetically distant obligate pathogens, we identify features of its signalling networks and secretome and we identify sequence features of its promoters. Finally, we present a set of microsatellite repeats that will be useful as markers for diversity and population studies.

## Resultsn

### General features of the genome assembly

We assembled Illumina HiSeq 2000 sequencing data from four genomic libraries into 2780 scaffolds. The draft assembly was relatively contiguous, with an N_50_ scaffold length of 246.53 kbp and an L_50_ scaffold count of 46 (Supplementary Table 1). The total length of 35.39 Mbp falls within the range observed for other *Peronospora* genome assemblies (Supplementary Table 1). In total, 9049 protein-coding genes were predicted from the repeat-masked genome. The circular mitochondrial genome was 40,106 bp long (Supplementary Figure 1).

Repeat sequences account for 5.71 Mbp, which is 16.15% of the total assembly size. Furthermore, 18.20 % of the scaffold-level assembly consisted of gaps between contigs. Nevertheless, more than 92% of the short-insert paired sequence reads were successfully mapped against the genome assembly, suggesting a high degree of completeness (Supplementary Table 2). This was also confirmed with BUSCO3 (Simão et al. 2015): of the 234 core genes of BUSCO’s Alveolata/Straminipila-specific library, 207 (88%) are found as intact and single-copy. This is similar to other *Peronospora* genome assemblies (Supplementary Figure 1), which contain between 94 (40%) and 221 (94%) complete single-copy genes. Of the core genes reported as missing by BUSCO3, most could be found by tBLASTNn searches, suggesting that the BUSCO3 underestimates the actual completeness of gene-space due to the divergence caused by high mutation rates in downy mildews (Thines and Kamoun 2010).

We note that the size of our genome assembly is smaller than that of a publically available, but unpublished *Pe. belbahrii* assembly (deposited by the University of Michigan, accession GCA_002864105.1), which is 59.2 Mbp in length and was assembled using Abyss2 (Jackman et al. 2017). However, BUSCO3 analysis reveals that the Michigan assembly contains a high frequency of gene duplication (Supplementary Figure 2). We also observed similar duplications when we assembled our data using Abyss2 instead of Velvet. The duplication level can be explained by assembly of distinct alleles into separate contigs. In line with this, the analysis of our genome data confirms substantial heterozygosity in the *Pe. belbahrii* genome (Figure 1).

**Figure 1.**
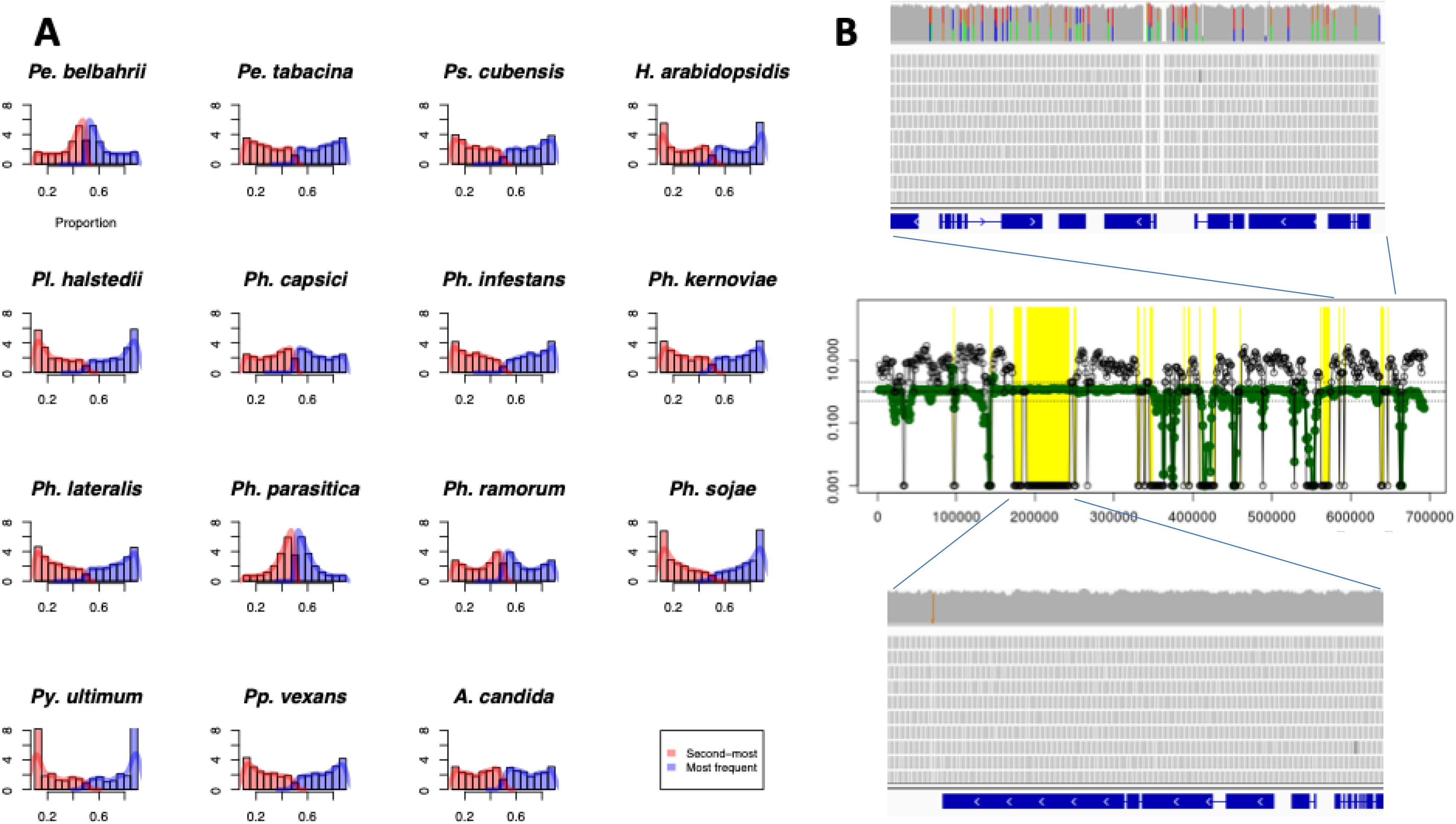
Heterozygosity. A: Frequency distributions of relative abundances of major allele over all genomic sites in sequenced oomycete genomes. For each species, raw Illumina sequence reads were aligned against the appropriate genome assembly using BWA-mem and discarding reads that align to more than one genomic location. For each single-nucleotide site in each genome, we counted the relative abundance of the most commonly occurring nucleotide (A, C, G or T). The histogram shows the frequency distribution for the relative abundance of the most commonly occurring nucleotide. The histograms have been cropped at the top for clarity. B: Distribution of heterozygosity across scaffold 882 of the *P. behlbarii* genome assembly. In the upper panel a heterozygous 72-kbp region is shown. Heterozygous sites are visible as two-coloured vertical lines, with the relative lengths indicating the relative abundance of the two bases in the reads aligned at that site. The central panel shows an overview of the heterozygosity profile and the depth of coverage (Y-axis) over the whole scaffold. The black circles indicate the number of heterozygous sites per 5-kbp window. The green circles indicate the normalised depth of coverage (divided by the coverage median, 51.6). The dashed horizontal lines indicate 0.5 x, 1 x and 2 x median depth of coverage. The yellow shading indicates regions where no heterozygosity was detected and depth of coverage was at least 90% of the median. The lower panel shows a 72-kbp region that lacks detectable heterozygosity.

## Heterozygosity

The genome of *Pe. belbahrii* displays significant levels of heterozygosity as indicated by a clear peak near to 50% allelic frequency in Figure 1A. A similar peak is also observed for some other oomycete genomes investigated (Supplementary Table 3), including those of Pe. *tabacina, Ph. ramorum* and Sa. *parasitica.* The distribution of heterozygosity over the *Pe. belbahrii* genome is not uniform and we identified several long stretches of the genome showing little or no heterozygosity. For instance, we identified a patchy distribution of heterozygosity over scaffold 882 (249 kbp), in which there is a region of 70 kbp that shows no detectable heterozygosity with normal sequencing depth, while other regions of this scaffold show more typical levels of heterozygosity as exemplified by another region of 72 kbp (Figure 1B).

### Gene spacing and promoter structure

Intergenic regions upstream of Pe. *belbahrii* start codons span a wide range of sizes (Supplementary Figure 3A), consistent with the organization of its genome into gene-dense (GDR) and gene-sparse regions (GSR); the former are defined as those in which genes have their 5’ ends within 2 kb of another gene. Seventy-six percent of predicted *Pe. belbahrii* genes are in GDRs, which is more than in other oomycetes such as *Ph. infestans* (51%) and *Pl. halstedii* (67%) (Haas et al. 2009; Baxter et al. 2010; Sharma et al. 2015). However, GSRs are likely to be disproportionately located in the unassembled gaps in draft-quality genome assemblies, meaning that these percentages should be interpreted with care. The median intergenic distance within *Pe. belbahrii* GDRs is 530 bp, which is slightly larger than in *Pl. halstedii* (420 bp) and *Ph. infestans* (430 bp).

*Peronospora belbahrii* promoters are relatively AT-rich, rising to 59% AT 50 bp upstream of the start codon (Supplementary Figure 3B). *Peronospora belbahrii* contains core promoter motifs at different frequencies as compared to other oomycetes (Supplementary Figure 3C). Each core motif occurs mostly within 100 bp of the start codon, as illustrated for the INR+FPR supra-motif in Supplementary Figure 3D.

As 64% of *Pe. belbahrii* promoters lacked a known core promoter sequence, attempts were made to identify alternative motifs using MEME (Bailey et al. 2009). Several candidates were identified, all of which resembled microsatellites. Figure 2A shows the most significant motif (p=10^-33^ compared to shuffled promoters), which is composed of TC dinucleotide repeats and showed a strong orientation bias (p=10^-40^; Figure 2B). A search of all *Pe. belbahrii* promoters for (TC)⊓ repeats with n=3 revealed significant over-representation of arrays having up to 8 repeat units (p<10^-3^ compared to shuffled sequences). A search for other dinucleotide microsatellites indicated that (TC/GA)_n_ was the most plentiful in *Pe. belbahrii,* with Z=5.7 (p<10^-6^) for over-representation compared to a shuffled dataset (Supplementary Figure 3C). Some other microsatellites were also overrepresented in promoters, including those with CT/AG, CA/TG, or AC/GT repeats.

**Figure 2.**
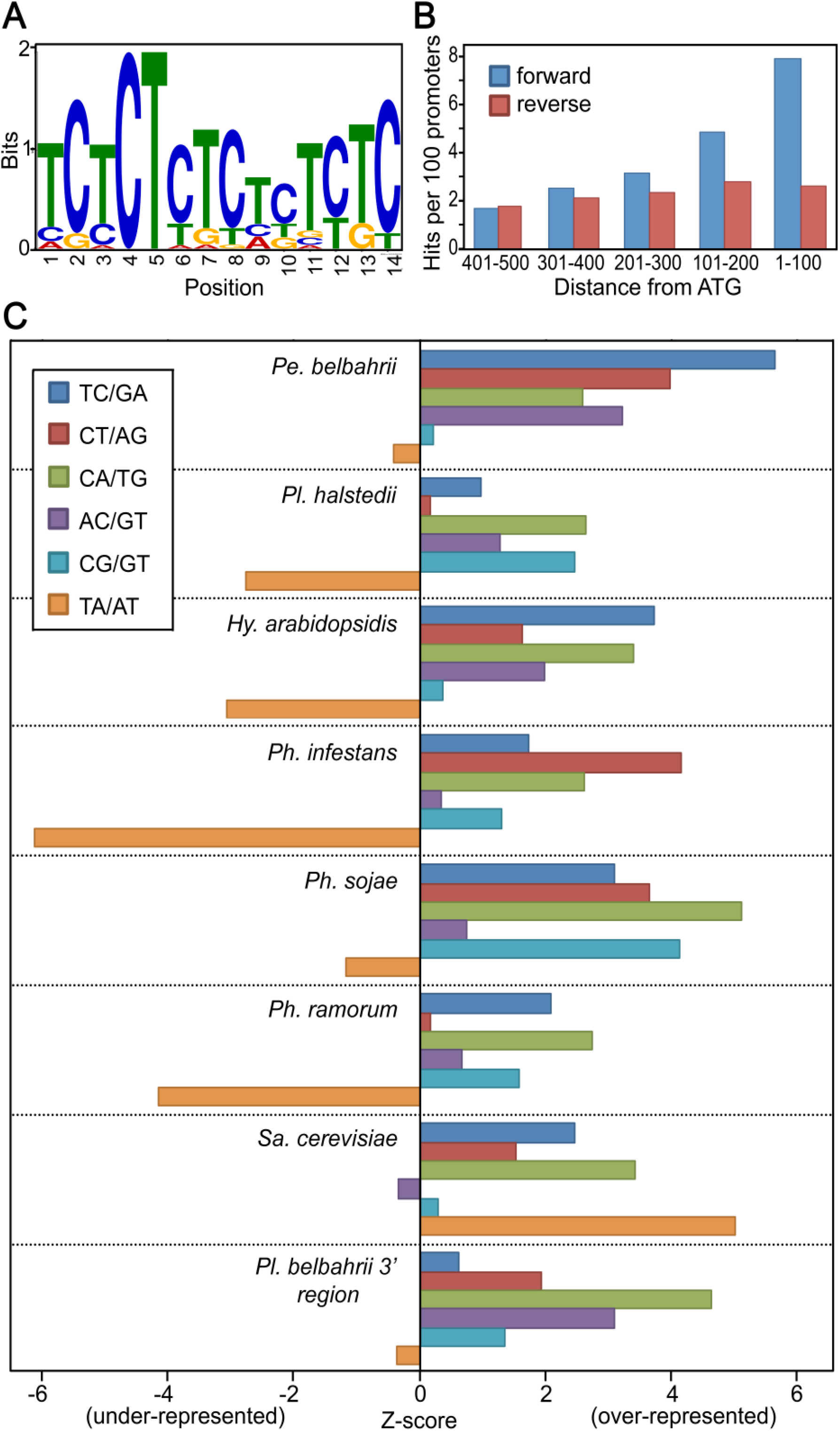
Microsatellite motifs in promoters. Panel A: TC-rich microsatellite-like motif detected in *Pe. belbahrii* promoters. Panel B: Orientation of the TC-rich motif in bins upstream of the start codon of *Pe. belbahrii* genes, based on matches with p<10^-4^. Panel C: Representation of microsatellites with 3 or more perfect repeat units in the 100 bp upstream of the start codons of *Pe. belbahrii, Pl. halstedii, Hy. arabidopsidis, Ph. infestans, Ph. soĵae, Ph. ramorum,* and *Saccharomyces cerevisiae,* and within 100 bp 3’ of *Pe. belbahrii* coding sequences. Z-scores (positive for over-representation, negative for under-representation) were calculated using a twice-shuffled promoter dataset as a control.

### Microsatellites

Microsatellites have potential use in identification processes and population studies. We characterized the full microsatellite complement of *Pe. Belbahrii* and compared it to 34 other species of pathogens. The proportion of microsatellite sequences in the majority of oomycete genomes ranged from 0.023% (*Al. candida*) to 0.331% (*Py. irregulare)* when only 2-6 bp arrays were considered. Higher ratios were observed for 1 to 6 bp simple sequence repeat (SSR) arrays (from 0.044% for *Pl. halstedii* to 0.341% for *Py. irregulare)* (Supplementary Tables 9-11). The fraction of the genome comprised of SSR arrays in fungi with respectively similar lifestyle was higher (0.146% for *Blumeria graminis* f. sp. *tritici* to 0.668% for *Colletotrichum graminifolia).* The proportion of SSRs in genomes was not correlated with the size of the assembled genomes (R^2^ = 0.0008). Interestingly, a low proportion of mononucleotides was observed in Sa. *parasitica* (less than 1%), *Pe. belbahrii* and *Pythium* species (3-10%). Dinucleotide and trinucleotide motifs were the most abundant types of repeats in the *Pe. belbahrii* genome, each representing more than 42% of all screened motifs (Supplementary Table 10). In coding regions of Pe. *belbahrii* di- and especially tri-nucleotide SSR arrays were frequently found, namely AAG/CTT, ATC/GAT an AG/CT (Supplementary Table 11). In oomycete genomes, a higher proportion of AAG/CTT and AGC/GCT among other 3 bp arrays was observed, while the presence of 4 to 6bp repeats was lower than in biotrophic fungi (Supplementary Tables 9 and 10). A total of 130 SSR markers (including 16 markers in coding regions) with potential use in population genetic studies were designed (Supplementary Table 12). These represent mainly di- and trinucleotide repeats (56 and 60 markers respectively), while markers with tetra- to hexanucleotide repeats are less abundant (4,1,1, respectively).

### Metabolism

To compare metabolism of the obligate biotroph Pe. *belbahrii* with that of other oomycetes, KEGG-based metabolic networks were constructed for 11 oomycete species with different lifestyles (Figure 2). Overall, the metabolic networks of the obligate biotrophs *Pe. belbahrii, Pl. halstedii, Al. laibachii,* and *Hy. arabidopsidis* contain fewer genes, enzymes, reactions and metabolites in comparison to the hemibiotrophs and the animal pathogen, Sa. *parasitica* (Figure 3). The enzymes in each metabolic network were associated with reactions of 108 different KEGG pathways (Supplementary Table 4). Species clustered together according to their lifestyle when classified by numbers of supported reactions per pathway (Figure 3), despite the fact that the non-peronosporalean *Al. candida* and *Al. labaichii* are phylogenetically not closely related to the downy mildews. As obligate biotrophy arose multiple times among the oomycetes, this suggests convergent evolution of metabolic functions and defects.

**Figure 3.**
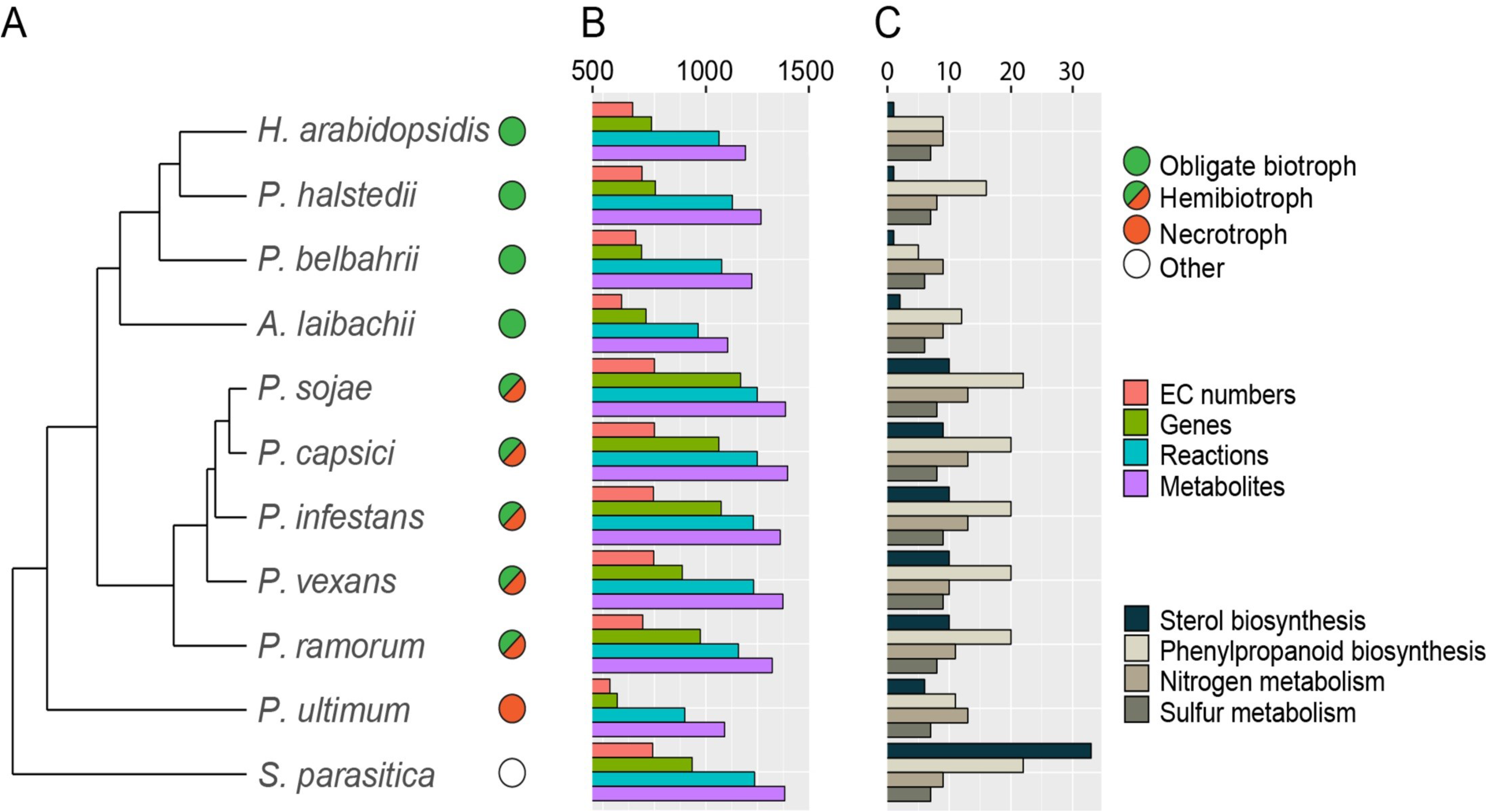
Metabolic pathway analyses in eleven oomycete species with different lifestyles (depicted by circles). (A) Hierarchical clustering of the species based on the number of reactions per KEGG pathway. (B) The number of different EC numbers, genes, reactions and metabolites included in the metabolic network and (C) the number of reactions identified in four metabolic pathways for each of the species.

Remarkably, the top ten most variant pathways (by coefficient of variation), contain three pathways classified in the KEGG BRITE category lipid metabolism (Supplementary Table 4). The steroid biosynthesis pathway was the most variable between species, in terms of supported reactions per pathway (Supplementary Table 4). *Peronospora belbahrii, Pl. halstedii* and *Hy. arabidopsidis* support only one reaction in this pathway, catalysed by a cholesterol ester acylhydrolase (EC: 3.1.1.13) and *Al. laibachii* supports one additional reaction, catalysed by an Acyl-CoA:cholesterol O-acyltransferase (EC: 2.3.1.26). The obligate biotrophs contain generally fewer enzymes that function in the phenylpropanoid biosynthesis pathway (KEGG: map00940) than other oomycetes. Interestingly, *Pe. belbahrii* is an extreme case supporting only five reactions in this pathway, whereas the number of reactions in other oomycetes ranges from 9 to 22. The analysis of primary metabolism pathways shows that all these mildews are unable to assimilate nitrate and nitrite and also lack an enzyme in the sulphur metabolism pathway generating cysteine (Figure 4), which is in line with the previous studies (Jiang et al. 2013; Baxter et al. 2010).

**Figure 4.**
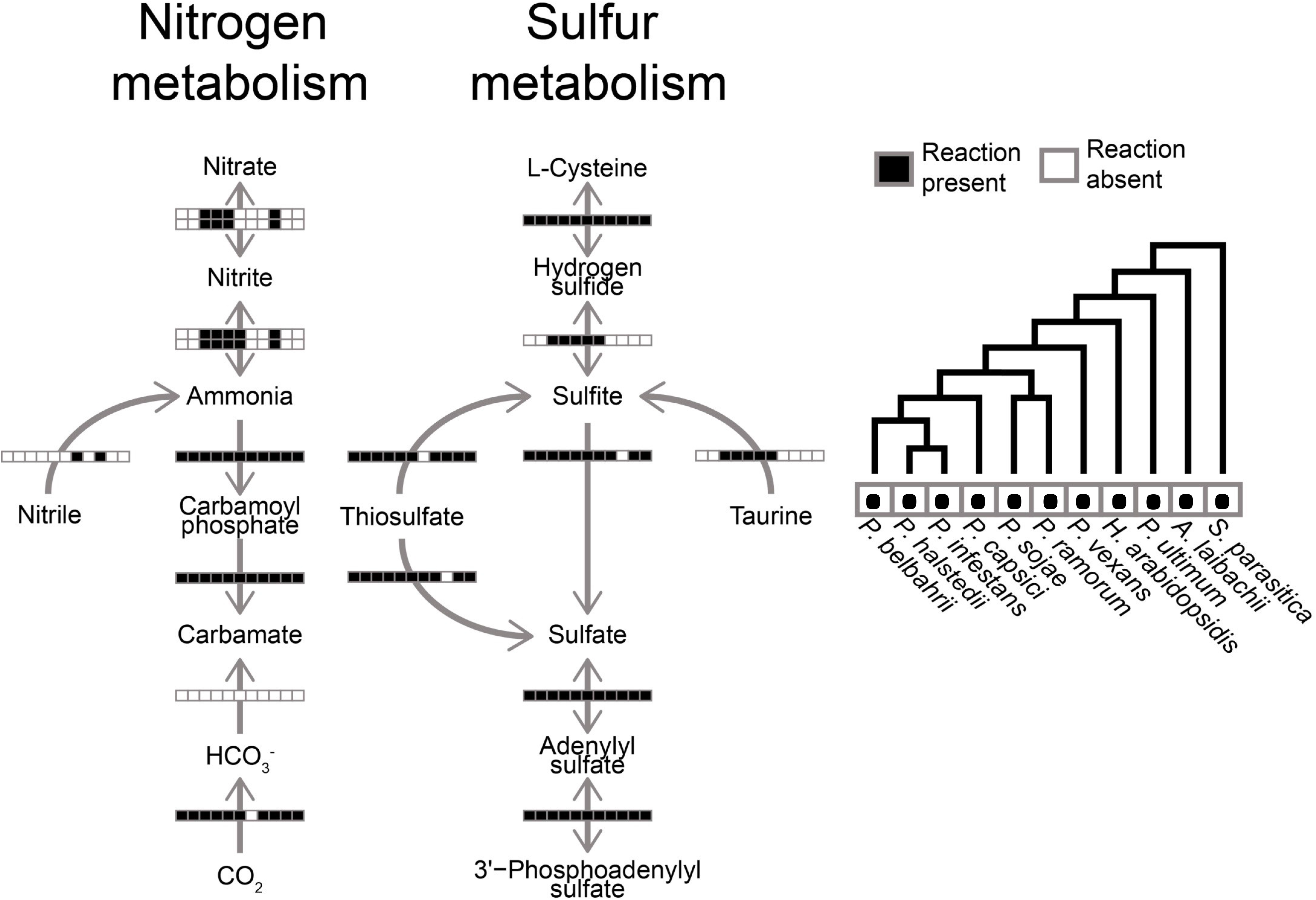
Presence/absence analysis of enzymes in the nitrogen and sulfur metabolism pathways. Arrows indicate the direction of the reaction. The horizontal bars are composed of black and white squares representing the species in the order of the phylogenetic tree on the right and presence/absence of the enzyme catalysing the reaction in the respective species.

In addition to the loss of metabolic pathways found in other oomycetes, *Pe. belbahrii,* like other members of the genus, does not produce zoospores. To check whether its genome has retained genes normally required for forming flagella, or remnants of such genes, sequences of 100 flagellum-associated proteins from *Ph. infestans* (Judelson et al. 2012) were used to search the *Pe. belbahrii* predicted proteome by using BLASTp and its genome using tBLASTn. No significant matches were detected, indicating a complete loss of its capacity to produce flagella.

### Signalling

Phospholipid modifying and signalling enzymes (PMSEs) may participate in both releasing carbon for metabolism and generating signalling molecules. Orthologs of nearly all genes encoding PMSE enzymes that have been identified in *Phytophthora* spp. and other oomycetes were found in *Pe. belbahrii* (Supplementary Table 5), but in comparison to *Phytophthora* spp. the total number is reduced (van den Hoogen et al. 2018). In *Pe. belbahrii,* as in *Hy. arabidopsidis* and *Pl. halstedii,* there are fewer genes coding for phospholipase D (PLD) compared to *Ph. infestans,* largely due to a decrease in sPLDs, the subclass comprising of PLDs that are likely secreted due to the presence of a signal peptide (Meijer et al. Govers 2011). Also type H of the phosphatidylinositol kinase (PIK) genes are missing. From the largest PMSE family in oomycetes, phosphatidylinositol phosphate kinases (PIPKs) with an N-terminal G-protein Coupled Receptor (GPCR) domain (GPCR-PIPKs), 11 members are detected in *Pe. belbahrii* (Supplementary Table 5). In common with other peronosporalean oomycetes, *Pe. belbahrii* lacks a canonical phospholipase C (PLC) gene. The only light-sensing proteins detected in the genomes of *Pe. belbahrii* and other oomycetes belong to the CRY/PHR superfamily. Each species contained three such proteins, which formed three well-supported clades in phylogenetic analyses (Supplementary Figure 4A). Each of the three *Pe. belbahrii* proteins (PBEL_00732, PBEL_04497, and PBEL_06190) had the canonical (Chaves et al. 2011) photolyase and FAD chromophore binding domains expected for CRY/PHRs (Supplementary Figure 4B). None had long C-terminal domains present on animal cryptochromes and are known to mediate circadian rhythm. *Peronospora belbahrii* also contained a fourth predicted protein (PBEL_00742), which was 99.4% identical at the nucleotide level to PBEL_06190, probably representing a duplicated gene.

### The secretome

Out of 9049 protein-coding genes, 413 were predicted as encoding secreted protein encoding genes. A set of 381 candidate genes encoding proteins with a leading signal peptide and no transmembrane domain was used as a starting point for further predictions and annotations. Additional curation resulted in 335 putative extracellular proteins (Table 1, Supplementary Table 6) organized as 76 tribes and 84 singletons. For roughly half of all proteins (171) it was possible to assign a particular functional category *(i.e.* not ‘hypothetical’). Annotated proteins with functions previously assigned to the infection process (effectors, pathogen associated molecular patterns (PAMPs), SCP, cell wall degrading enzymes, proteases) comprise 31% of all secreted proteins (Table 1). In general, when compared to other obligate parasitic oomycetes, we found reduced repertoires of secreted proteins in the *Pe. belbahrii* secretome (Table 1). Strikingly, we identified no CRNs with predicted signal peptides and only two NPP-like proteins (NLPs). Proteins with similarity to CRNs but lacking a predicted signal peptide have been found in oomycete genomes previously, but the low number of NLPs is unprecedented.

**Table 1.**
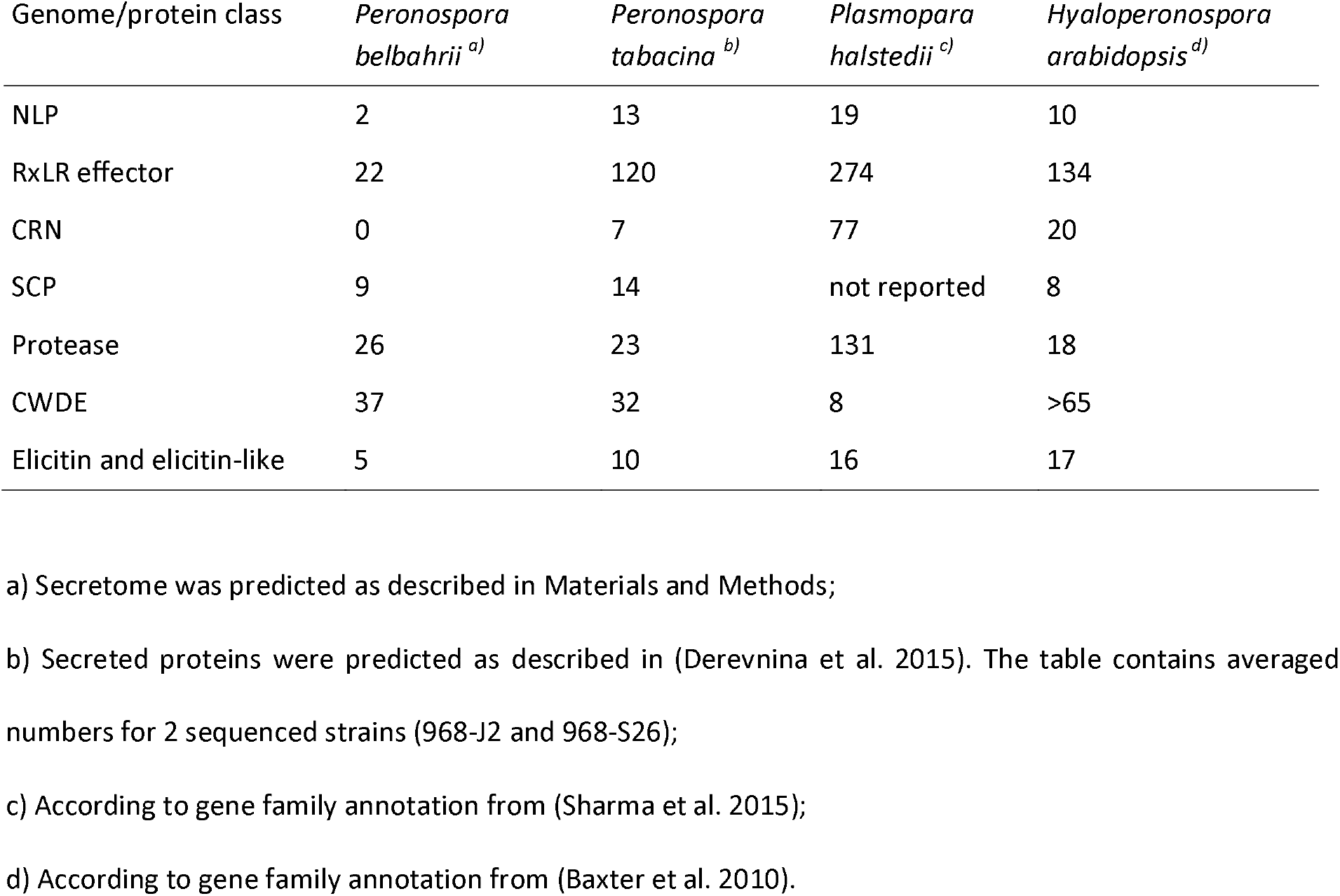
Classes of secreted proteins in selected oomycete genomes.

Among predicted effectors, the most abundant category encompasses RxLR-like effectors (22 proteins), in three of which (PBEL_01290, PBEL_01294, PBEL_02639) share sequence homology with known oomycete avirulence genes from *Ph. sojae* (Avh154, Avh160 and Avh152). Of 22 predicted RXLR effectors, 14 have the exact RxLR motif, while 5 harbour a RxLK motif, and three singletons have other variations of the motif (Figure 5).

**Figure 5.**
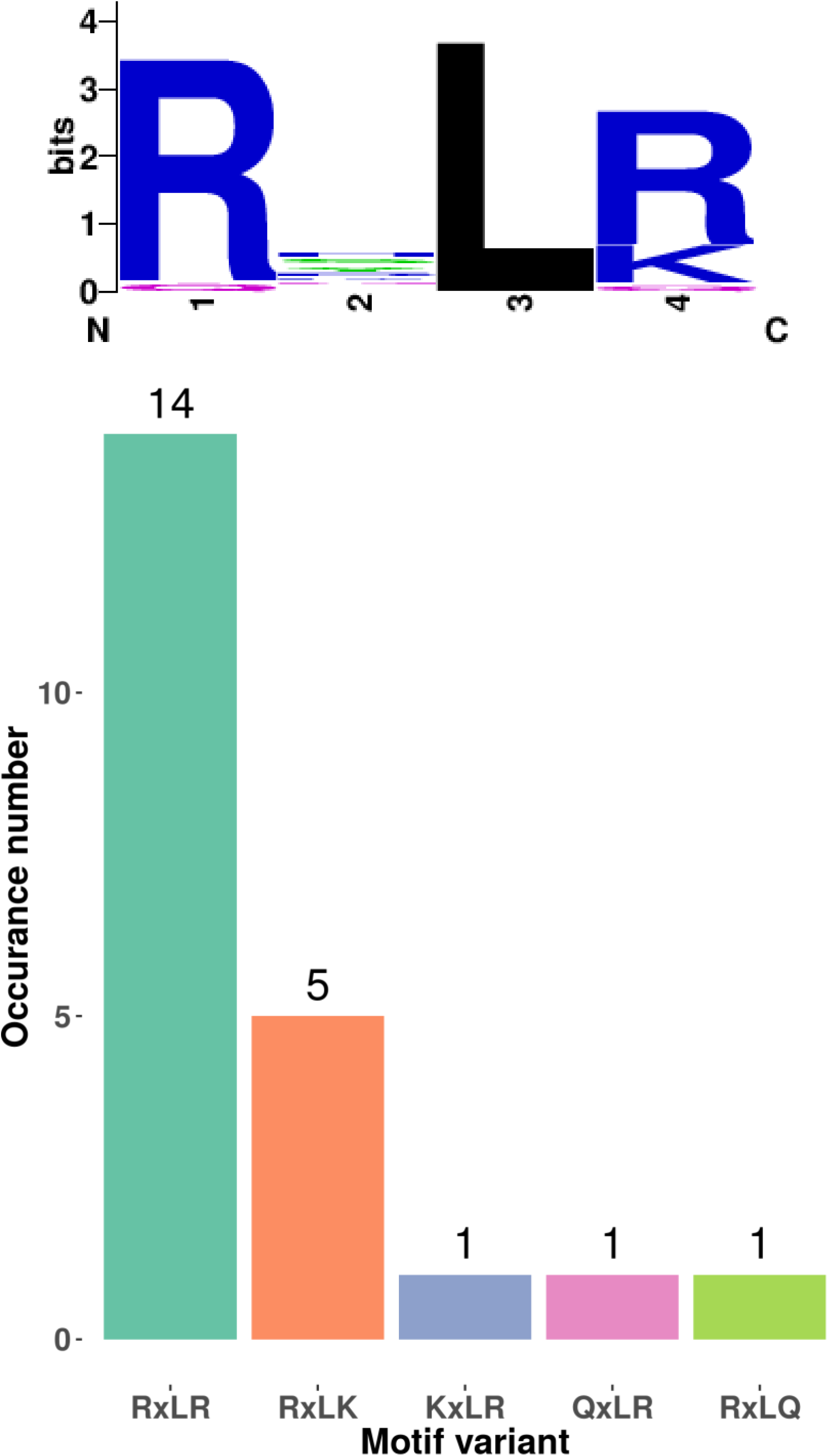
RxLR motif conservation. Sequence Logo for prediceted RxLR-like effectors of *Pe. belbahrii.* Only 14 canonical RxLR effectors and 8 variants are present in the predicted secretome, which is by far lowest number for any *Phytophthora* or downy mildew genome sequenced so far.

Only two protease inhibitors were identified (PBEL_30414, PBEL_30415), both having sequence similarity to Kazal-like serine protease inhibitors (Supplementary Tables 6 and 7, Table 1).

Six *Pe. belbahrii* predicted secreted proteins contain potential NLS signals (Supplementary Table 7). Presence of an NLS in combination with a signal peptide suggests a role of these proteins in the host nucleus, yet only one (PBEL_07289) carries a recognisable RxLR motif implied in host-translocation, while six other (PBEL_06934.2, PBEL_06683, PBEL_07130, PBEL_04801, PBEL_08621.2 and PBEL_05668) might remain extracellular or be translocated by an RxLR-independent mechanism.

The *Pe. belbahrii* secretome contains several hydrolytic enzymes (63, 19% of all extracellular proteins), which we classified into two main categories: cell wall degrading enzymes (CWDE) and proteases. CWDE are mainly represented by glycosyl hydrolases (24), while proteases are dominated by serine and cysteine proteases (Table 1). We detected only five proteins with similarity to known immune response elicitors, represented exclusively by elicitins.

To identify common and specific secreted protein families, OrthoMCL was used to group predicted secretome members of *Pe. belbahrii* with those from several plant pathogenic oomycetes into protein families based on homology. Eleven families representing 23 proteins solely contain sequences from *Pe. belbahrii.* Nine out of the 23 proteins (39%) are predicted as RxLR effectors whereas the predicted RxLR effector proteins only make up 7.4 % of the total predicted secretome. Furthermore, 91 *Pe. belbahrii* proteins did not cluster with any other protein in the set and are probably unique to the *Pe. belbahrii* secretome (Supplementary Tables 6 and 7).

Twenty-three ortholog families with a combined number of 68 genes were found to be exclusively present in one or more downy mildews. Of these, 12 families (35 genes) are not shared with *Pe. belbahrii.* A BLASTP search of the other families against the NCBI non-redundant database on the full proteins (including signal peptide) showed that the best non-downy mildew matches for the *Pe. belbahrii* proteins in six of these families do not have signal peptide (Supplementary Table 8). These proteins have potentially evolved to acquire a signal peptide and a new function in downy mildews.

### Discussion

We assembled and annotated the genome sequence for *Pe. belbahrii* with the aim to produce a useful resource to the community for follow-up studies. The size and quality of the resulting genome are comparable to genome assemblies of other members of the genus and other oomycetes. While the genome is similar to many other oomycetes in terms of size and architecture, a remarkable diversity in the occurrence of dinucleotide repeats in promoters was observed. An analysis of other oomycetes revealed remarkable diversity in such microsatellites (SSRs). The (TC/GA)_n_ repeats were over-represented strongly not only in promoters *Pe. belbahrii* but also in *Hy. arabidopsidis* and three species of *Phytophthora,* but not in *Pl. halstedii.* Notably, (TC/GA)_n_ elements were not overrepresented in the regions downstream of *Pe. belbahrii* genes. Interestingly, (TA/AT)n showed a strong tendency towards under-representation in most oomycete promoters, even though it is over-represented in members of several other eukaryotic kingdoms including fungi (*Saccharomyces cerevisiae)* (Bunnik et al. 2014; Sawaya et al. 2013; Vinces et al. 2009). In other species, dinucleotide repeats are believed to affect the positioning of nucleosomes on DNA (Bunnik et al. 2014; Sawaya et al. 2013; Vinces et al. 2009). Tandemly repeated DNA sequences in promoters are of interest since they display a propensity to mutate, which may enable the tuning of gene expression by affecting chromatin structure (Vinces et al. 2009). Underrepresentation of the AT-rich repeats in most oomycetes is in itself likely a signal of function, and is suggestive of divergence in organization of chromatin in oomycetes. In many cases, SSR repeat number appears to be a key factor that determines gene expression and expression level. Some genes can only be expressed at a specific repeat number of SSRs. For example, the GAA array, which is the most frequent trinucleotide repeat *(i.e.* AAG/CTT) in the majority of analysed oomycete genomes including *Pe. belbahrii,* is reported to be the key factor influencing the promoter activity of the *Escherichia coli lacZ* gene (Liu et al. 2000). In addition to their potential importance in regulating gene expression, SSRs are also of considerable interest for population genetics studies. Therefore, a set of 130 microsatellite markers was developed in this study, which may prove to be a useful tool for future population genetic studies in *Pe. belbahrii* and closely related species on other culinary herbs.

So far, relatively little is known about oomycete metabolism. Several metabolic pathways have been affected by the consequences of the close symbiosis of downy mildews and their hosts, which led to a reliance on the host metabolism to provide nutrients for the pathogen (Thines 2014; Judelson 2017). It is interesting to note that based on hierarchical clustering of metabolic networks the different oomycete species clustered together according to their lifestyle, and that therefore *Albugo candida* clustered with the downy mildews, irrespective of its distant phylogenetic relationship to them (Thines 2014). Similarly, the hemibiotrophic *Phytophthora* species clustered together, forming a sister-group to the obligate biotrophic taxa. This highlights that similar evolutionary forces due to the adaptation to similar lifestyles will lead to convergent patterns (Kemen et al. 2011; Sharma et al. 2015b). In line with previous studies, we found that the metabolic networks of obligate pathogens are generally smaller (Rodenburg et al. 2018; Spanu 2012) and lack enzyme-encoding genes for reactions in various metabolic pathways (Judelson 2017; Kemen et al. 2011; Sharma et al. 2015a; Baxter et al. 2010).

Many cellular pathways are linked to phospholipid signalling, including not only metabolic pathways, but also pathways regulating the expression of genes required for successful colonisation. Despite the importance of phospholipid signalling demonstrated in many organisms, in oomycetes only a few PMSEs have been studied in detail. The domain composition of GPCR-PIPKs, shared by all oomycetes, points to a mechanism that directly links G-protein mediated signalling with phospholipid signalling (van den Hoogen and Govers 2018). Silencing of a PIK-A and PIK-D in *Ph. soĵae* showed that these two PIKs are required for full virulence (Lu et al. 2013), and GPCR-PIPKs were found to be involved in sporangia development, chemotaxis, zoospore development, and oospore development (Hua et al. 2013; Yang et al. 2013). Given this wide span of lifecycle stages affected, it seems possible that also other developmental stages are influenced by phospholipid signalling, such as formation of the hyphal network and the production of haustoria.

Extracellular effector proteins secreted from hyphae and haustoria play key roles in infection strategies of pathogenic oomycetes helping to establish colonization and to modulate plant responses (Meijer et al. 2014). By investigating the protein domains of secreted proteins their role in infection can be clarified. While a large part of the secretome of Pe. *belbahrii* could be classified through sequence similarity, 203 proteins remain hypothetical and could represent yet-to-be discovered effector proteins.

Besides the canonical effector proteins such as RXLRs, two proteins that possess a carbonic anhydrase domain (Pfam: PF00194) were identified in the secretome. In a study of the secretome of *Phytophthora infestans,* carbonic anhydrases were found to be expressed in planta during infection of potato and they were suggested to be potential new virulence factors (Raffaele et al. 2010). In addition, the secretome comparison revealed 23 families, containing 68 proteins, that are only found in downy mildews but not in more distantly related *Phytophthora, Pythium* or *Saprolegnia* species. Of these, 10 families (28 proteins) include proteins derived from *Pe. belbahrii.*

Remarkably, a similarity searches BLASTp revealed that proteins in six of these families had an ortholog without signal peptide in other oomycetes, like in *Phytophthora* species. Possibly these genes are present in other oomycetes and have evolved to encode for secreted proteins within the lineage of the downy mildews. Some of these proteins have interesting domains, *e.g.* DUF953 (Seidl et al. 2011), E3 ubiquitin-protein ligase (Zeng, et al. 2006), or bZIP_2 (Gamboa-Meléndez et al. 2013). For proteins in the last two downy mildew families that have a non-downy mildew ortholog without a predicted signal peptide, no known domain was identified. Interestingly, one of these families is the largest family only present in downy mildews, encompassing five proteins from three different species *(Pe. tabacina, Pe. belbarhii* and *Pl. halstedii).* The other group has three proteins from three species *(Pe. belbarhii, Hy. arabidopsidis* and *Pl. halstedii.* The functions of these proteins are unknown and since they seem to be present in the secretome of downy mildew species but not of other oomycetes included, they are interesting candidates for functional validation.

Strikingly, we found an underrepresentation of two common groups of effectors, the CRNs and NLPs compared to other secretomes. It is possible that CRNs are underrepresented as they might have not been predicted by SignalP, for several known CRNs have non-canonical signal peptides (Stam et al. 2013). However, the generally low number of classical effectors, such as the potentially necrosis-causing CRNs, NLPs, and RxLRs in *Pe. belbahrii* might reflect the combined effects of adaption to biotrophy and host specificity.

Of the only 22 RxLR-like effectors found in *Pe. belbahrii,* seven did not have orthologs in other species Possibly, these effectors can be considered species-specific or have become too dissimilar from their orthologs to be recognized. Among the RxLR effectors, PBEL_07289 is a notable member, with C-terminal homology to Nudix hydrolases. The effector AVR3b *Ph. soĵae,* another NUDIX-domain containing RxLR effector has been demonstrated to increase susceptibility to *Ph. capsici* and *Ph. parasitica* (Dong, et al. 2011). It is noteworthy that the genome of *Pe. belbahrii* was predicted to contain the lowest number of RxLR-like effectors compared to all other sequenced Peronosporaceae. This might hint at a very high specialisation of *Peronospora* species (Voglmayr 2003; Thines et al. 2009; Choi et al. 2009), necessitating only few highly effective effectors, as targets need to be manipulated mostly in just one or very few closely related host species (Thines and Kamoun 2010). Alternatively or in addition, this might reflect the evolution of few highly efficient core effectors that also enable *Pe. belbahrii* to extend its range (Thines 2019) and affect other related species under optimal conditions (Naim et al. 2019).

In conclusion, analyses of the *Pe. belbahrii* genome revealed a rather small oomycete genome with little more than 300 secreted proteins. We found a remarkable signal of convergent evolution with respect to lifestyle in various oomycete genomes with metabolic pathway losses and patterns of diversification. Our analyses suggest a divergent organisation of promoters, as suggested by the high frequency of CT-repeats. In addition, we found the lowest amount of canonical effectors in any oomycete genome sequenced so far, but found hints for the presence of virulence factors that have so far not been investigated, and which are promising candidates for future functional studies.

## Materials and methods

### DNA isolation from spores and library preparation for genomic sequencing

Several heavily infected basil plants with fresh sporulation (violet-brown down on the lower leaf surfaces) from a single lot were bought from a supermarket in Butzbach, Germany in summer 2015. From these, several leaves were rinsed with deionised water to collect spores. DNA was extracted from spores pelleted by centrifugation as described previously (Sharma et al. 2015a). RNA was extracted from fresh leaves with freshly sporulating pathogen as described previously (Sharma et al. 2015). DNA and RNA extracts were sent to a commercial sequencing provider (LGC Genomics, Germany) for preparation and sequencing of 300 bp, 800 bp, 3 kbp, and 8 kbp genomic libraries and a 300 b.p. library for RNA-Seq. Paired-end Illumina HiSeq sequencing was carried out with 100 bases sequenced from both ends of the fragments in the libraries.

### NGS data processing and genome assembly

Standard primers and adapter sequences were clipped from the raw genomic Illumina reads using Trimmomatic (Bolger et al. 2014). A window-based quality filtering using Trimmomatic was performed by keeping an average quality cut-off of 20 within a window of size 5 bp. Reads shorter than 65 bp were filtered out. The second round of filtering was performed using FastQFS (Sharma and Thines 2015). Read pairs were eliminated if either read had an ambiguous base, or if an individual base had a Phred-scale quality score of 3 or lower, or if the average Phred score was below 26. The filtered reads were assembled using Velvet v1.2.09 (Zerbino and Birney 2008). Several different assemblies were generated using different k-mer sizes and other parameters, including expected coverage and coverage cut-offs. Different assemblies were compared and the best assembly was decided on the basis of N_50_, L_50_, number of scaffolds and genome completeness using Quast (Gurevich et al. 2013) and BUSCO3 (Simão et al. 2015). Assemblies were also generated using SPAdes version 3.12.0 and SOAPdenovo version 2.04, exploring a range of k-mer sizes; however, these yielded less-complete and less-contiguous assemblies than the Velvet assembly. The mitochondrial genome was assembled from the Illumina short reads, and a mitochondrial backbone sequence of *Pe. tabacina* (Derevnina et al. 2015) using the Mitomaker pipeline (available at: http://sourceforge.net/projects/mitomaker/). The physical map of the genome was visualised using OGdraw (Lohse et al.2007).

### Repeat element and gene prediction

Repeat elements were predicted and masked using the RepeatScout v1 (Price et al. 2005), RepeatModeler v1.0.4 (http://www.repeatmasker.org/RepeatModeler.html) and RepeatMasker (Tarailo-Graovac and Chen 2009) pipelines as described before (Sharma et al. 2015). The repeat-masked scaffolds were used for gene predictions. Both *ab initio* and transcript-based gene prediction tools were used to define gene boundaries as described previously (Sharma et al. 2015) (Supplementary Figure 5). Translated protein sequences were searched for an extracellular secretion signal and transmembrane domain using SignalP (Petersen et al. 2011), TargetP (Emanuelsson et al. 2000) and TMHMM (Krogh et al. 2001) as described (Sharma et al. 2015). Due to low RNA-Seq support for some genes predicted to code for secreted proteins, gene boundaries of potential secreted effectors were manually curated by looking for potential alternate start codons up to 280 bp upstream of the start codon predicted by evidence modelling.

### Estimation of heterozygosity

The availability of shotgun sequencing reads at high coverage, drawn randomly from both chromosomes from each homologous pair, offers the opportunity to estimate patterns of heterozygosity across the genome of various oomycetes (Supplementary Table 3). First, poor-quality sequence read pairs were removed using TrimGalore (http://www.bioinformatics.babraham.ac.uk/projects/trim_galore/) with the −q 30 and −paired options and then the remaining read pairs from the 300 bp library were aligned against the *Pe. belbahrii* genome assembly using BWA-mem (Li and Durbin 2009). Multi-mapping reads were eliminated using samtools view (Li et al. 2009) with the −q 1 option. For each single-nucleotide position in this alignment of reads against the genome assembly, we counted the frequency of each of the four possible bases in the reads aligned at that site. Thereby, for each site in the genome, we were able to estimate whether it was likely to be homozygous (most common base close to 100% frequency and second-most common close to zero) or heterozygous (close to 50% most common base and for second-most common base). Using R (R Development Core Team 2013), we plotted a histogram describing the frequency distribution of relative abundance of the most abundant base and second-most abundant base at each position in the genome. If the genome is highly homozygous, then the histogram would be expected to display a single peak close to 100% abundance for the most common base and a single peak close to zero for the second-most abundant. However, heterozygous sites would contribute to a second peak close to 50% abundance of the most common base and for the second-most abundant base. We also took a sliding-window approach to identify regions of high and low heterozygosity relative to the average over the whole genome. Each non-overlapping window of 5 kbp was examined for heterozygous sites, *i.e.* sites where the abundance of the most common base was between 48 and 52%. The density of heterozygous sites was defined as the number of heterozygous sites per 5-kb window. Density of heterozygous sites was plotted along with depth of coverage against genomic position, using R. The R scripts used for these analyses are available from https://github.com/davidijstudholme/heterozygosity.

### Metabolic networks

Metabolic networks were reconstructed for *Pe. belbahrii, Pl. halstedii, Ph. infestans, Ph. capsici, Ph. soĵae, Ph. ramorum, Phytopythium vexans, Hy. arabidopsidis, Py. ultimum, Albugo laibachii* and *Saprolegnia parasitica* using the RAVEN toolbox (Agren et al. 2013). Briefly, KEGG orthologous (KO) groups of enzymes (Chen et al. 2016; Kanehisa et al. 2012) were aligned using ClustalW2 v2.0.10 (Larkin et al. 2007), and based on these multiple sequence alignments, hidden Markov models (HMMs) were trained using HMMER v2.3 (Eddy 2011). Subsequently, the proteomes were matched to the HMMs with an E-value threshold of 10^-50^. The resulting enzyme orthologs were associated to KEGG reactions, which are linked to compounds and pathways. A metabolic reaction may occur if one or more catalysing enzymes are encoded in the genome. Hence, for each KEGG metabolic pathway, the number of supported reactions per species was counted. For these numbers, the coefficient of variation (variance/mean) was used to select the most variant pathways between species. In addition, these numbers were used to perform hierarchical clustering (UPGMA) of the species, using 1 minus the Spearman correlation coefficient as a distance measure.

### Promoter structure

Core promoter motifs as defined for *Ph. infestans* (Roy et al. 2013) were used to search 200 b.p. of DNA 5’ of *Pe. belbahrii* coding sequences using FIMO (Bailey et al. 2009). Searches for new motifs were performed using MEME (Bailey et al. 2009). Microsatellites within promoters were scored using MICAS (Sreenu et al. 2003). Twice-shuffled promoter sequences were used as control datasets, and Fisher’s Exact Test was used for tests of significance.

### Microsatellite motifs and prediction of SSR markers

The overall statistics of the most frequent motif types, repeat numbers and length of microsatellite sequences in *Pe. belbahrii* genome were calculated in Phobos (Mayer et al. 2010). SSRs with perfect repeat motifs ranging from mono- to hexa-nucleotides were considered, with the minimum number of ten (1 bp) or five (2-6 bp) repeat units. Searches of each motif were performed in separate runs *(i.e.* for 3 bp array the “Repeat unit size range” was set “from 3 to 3” and “Minimum length” set to “15 + 0 but not less than 15”), with results exported in “one-per-line” format, “Printing mode” set to “normalized – cyclic + rev. complement”, and all other settings set as default. Subsequent data analyses were conducted in Microsoft Excel. For comparison, the genomes of other oomycetes and selected fungi (Supplementary Table 13) were analysed as for *Pe. belbahrii.*

To facilitate the design of *Pe. belbahrii* SSR markers, only perfect di-to hexanucleotide repeats, with minimum repeats set to 8, 7, 6, 6, and 6 respectively, were extracted using Phobos. Primer pairs were designed using Primer3 (Untergasser et al. 2012) yielding PCR products of 150–500 bp. Data resulting from SSR analyses refer to duplex DNA, even if we show only the sequence of the repeated motif on one strand for simplicity, *i.e.* notations like AC and (AC)_n_•(GT)_n_ are equivalent. To see if the proportion of SSR arrays is correlated with the size of an assembled genome, linear regression in MS Excel was performed. Finally, the relative abundance (calculated as the ratio of the number of microsatellites per Mb of sequence analysed) and relative density (the ratio of the microsatellite length in bp per Mbp of the sequence analysed) of microsatellites were calculated separately.

### Light sensing and phospholipid signalling

Representative light-sensing proteins from other eukaryotes were used to query a database of predicted proteins from *Pe. belbahrii, Pl. halstedii, Hy. arabidopsidis,* and *Ph. infestans* using BLASTP. If no hits were found, the entire genome was searched using tBLASTn. Protein alignments were made using MUSCLE and trees generated using PhyML using the LG distance model.

The genome of *Pe. belbahrii* was screened for phospholipid modifying and signalling enzymes (PMSE)-encoding genes. A database of *Ph. infestans, Hy. arabidopsidis* and *Pl. halstedii* PMSE protein sequences was created and BLASTP and tBLASTn (Altschul et al. 1990) and searches were performed with an e-value cut-off score of le^-10^. Hits were manually inspected and PMSE gene models were linked to gene IDs. All gene models were manually inspected. Multiple sequence alignments on catalytic domains were made using MUSCLE (Edgar 2004), iterated 100 times, and phylogenetic trees were built using Neighbor-Joining with the Jukes-Cantor genetic distance model in Geneious R9 (Kearse et al. 2012).

### The secretome

For annotating candidate secreted proteins, profiles were built using several approaches: BLASTP search against GenBank NR database with E-value cut-off of 10^-6^; InterProScan v5.16 (Mulder et al. 2005) search against databases of functional domains with default parameters; RXLR and EER motif prediction using regular expressions; WY motif prediction based on WY-fold HMM using the hmmsearch function in the HMMR3 package (http://hmmer.org/); LxLFLAK and HVLVVVP motif predictions based on an HMM model built using known CRN effectors from other oomycetes according to Haas et al. (2009); and NLS motif prediction by NLStradamus version 1.8 (Nguyen Ba et al. 2009), posterior threshold = 0.6 and PredictNLS (Cokol et al. 2000) with default parameters. All obtained data are aggregated in Supplementary Table 6. Functional categories were assigned based on manual curation of the resulting table. The category ‘Hypothetical’ was assigned to proteins either having similarity to only hypothetical proteins or when the best 20 hits of the BLASTP output did not show consistency in terms of distinct functional categories. Proteins having significant sequence similarity to ribosomal, transmembrane proteins or proteins with known intracellular localization *(e.g.* heat shock proteins) and/or having respective domains identified by InterProScan were marked as possible false predictions. The contamination category was assigned for proteins with significant sequence similarity (revealed by BLASTP) to amino acid sequences from phylogenetically distant taxa *(e.g.* plants or mammals). It should be noted that this category will also include highly conserved proteins. Entries marked as both ‘false prediction’ or ‘contamination’ were excluded from further consideration.

Nine oomycete genomes were selected for secretome comparison: three downy mildew species (*Hy. arabidopsidis, Pe. tabacina* (two strains), and *Plasmopara halstedii),* three representative *Phytophthora* species *(Ph. infestans, Ph. soĵae* and *Ph. ramorum),* the more distantly related plant pathogen *Pythium ultimum,* and the fish-parasitizing oomycete *Saprolegnia diclina.* For each of these species the set of all predicted proteins was used to filter out the proteins that belong to the secretome as described for the *Pe. belbahrii* secretome. OrthoMCL 2.0 (Fischer et al. 2011) was used to create groups of orthologous proteins, with default settings.

## Supporting information

All 19 supplementary files

## Acknowledgements

This work was supported by the research funding program LOEWE “Landes-Offensive zur Entwicklung Wissenschaftlich-ökonomischer Exzellenz” of Hesse’s Ministry of Higher Education, Research, and the Arts in the framework of IPF and the LOEWE Centre for Translational Biodiversity Genomics (TBG). Further funding was received by the Food-for-Thought campaign from the Wageningen University Fund (SYAR, FG), the Research Council Earth and Life Sciences (ALW) of the Netherlands Organization of Scientific Research (NWO) in the framework of a VENI grant (863.15.005) (MFS) and a JSTP grant (833.13.002) (DJvdH, FG), the National Institute of Food and Agriculture and National Science Foundation of the United States (HSJ, the Gatsby Charitable Foundation (GAT3395/GLD)(AG, SS) and the Royal Society (UF160413) (SS).

## Supplementary Files

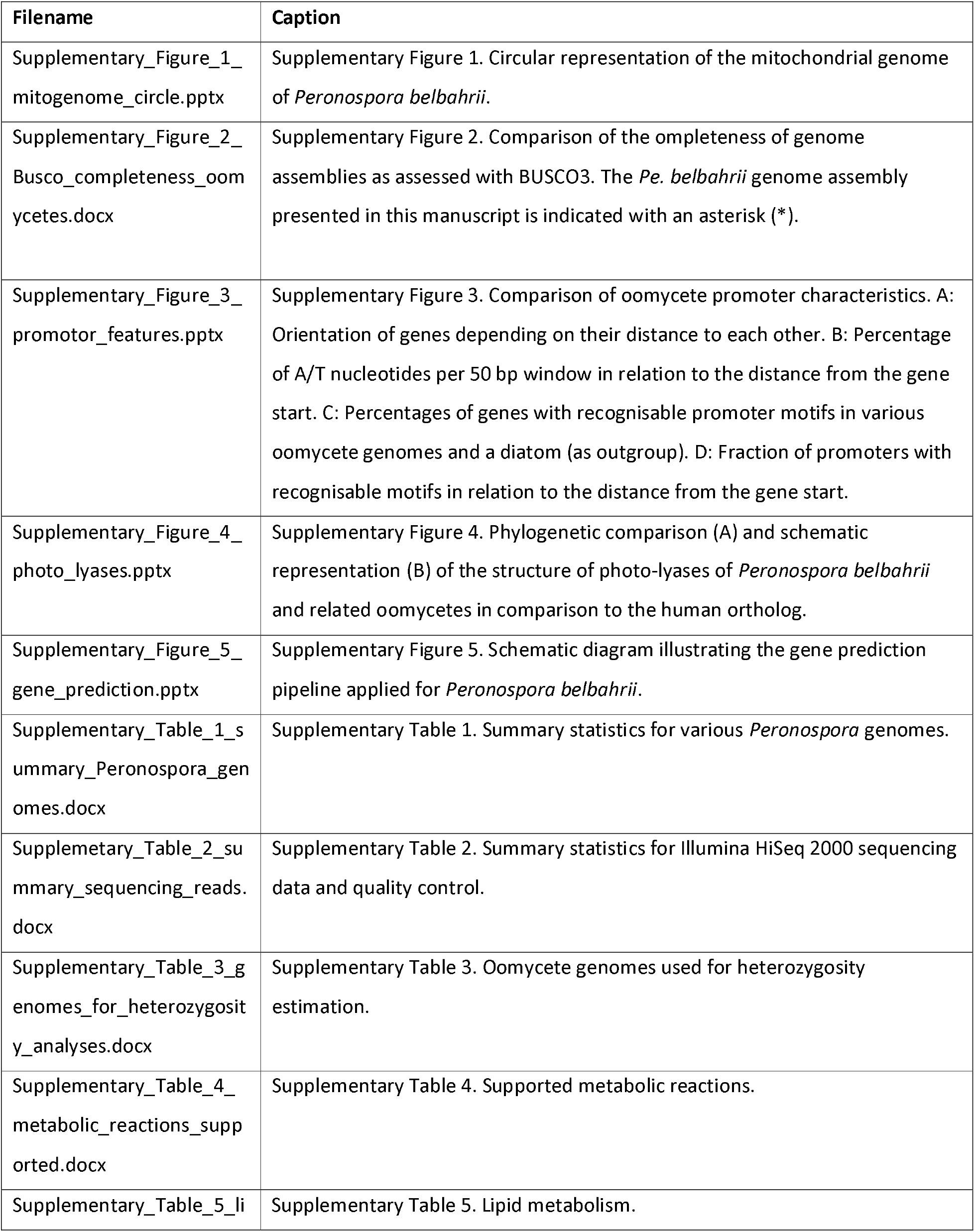
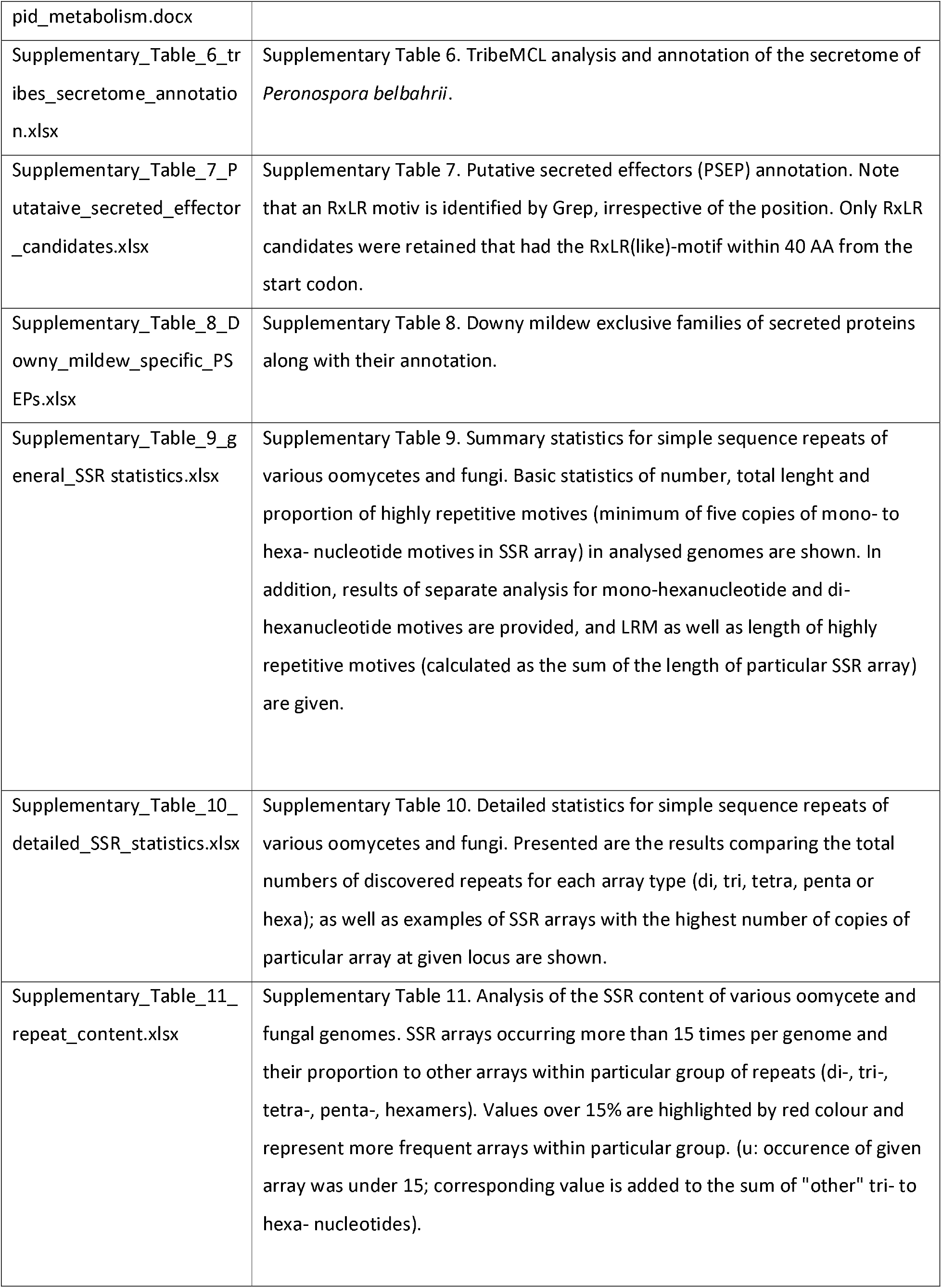
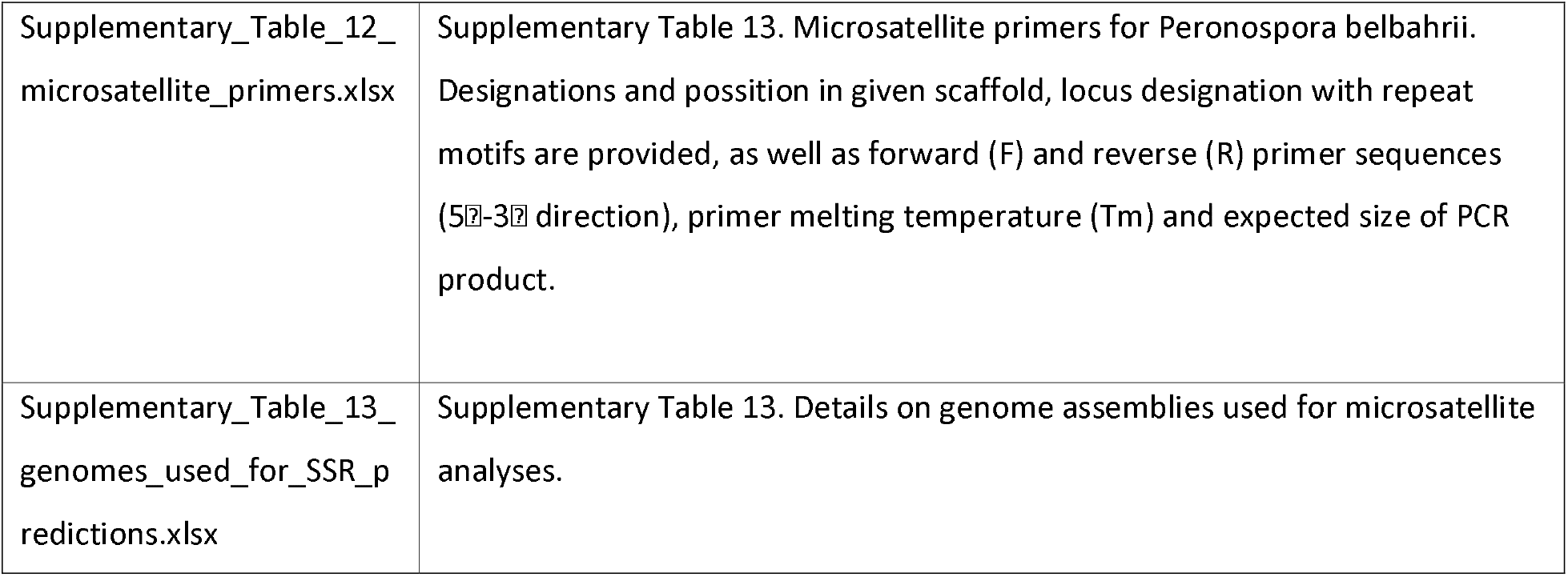

