## Supplementary material for "The genome of *Peronospora belbahrii* reveals high heterozygosity, a low number of canonical effectors and CT-rich promoters": All 19 supplementary files: Supplementary_Figure_2_Busco_completeness_oomycetes.docx

|  | Complete, single-copy | Complete, duplicated | Fragmented | Missing |
| --- | --- | --- | --- | --- |
| Pythium ultimum ADOS01000001.1 | 226 | 1 | 2 | 5 |
| Phytophthora sojae GCA_000149755.2 | 222 | 7 | 1 | 4 |
| Plasmopara muralis GCA_003676415.1 | 222 | 0 | 2 | 10 |
| Peronospora effusa GCA_002245715.1 | 221 | 0 | 2 | 11 |
| Peronospora effusa GCA_003704535.1 | 221 | 0 | 0 | 13 |
| Peronospora effusa GCA_003843895.1 | 221 | 1 | 2 | 10 |
| Peronospora effusa GCA_002245725.1 | 220 | 0 | 2 | 12 |
| Peronospora effusa GCA_002245735.1 | 220 | 0 | 2 | 12 |
| Plamopara viticola GCA_001695595.3 | 219 | 4 | 1 | 10 |
| Phytophthora infestans GCA_000142945.1 | 218 | 6 | 1 | 9 |
| Phytophthora ramorum AAQX01000001.1 | 218 | 8 | 3 | 5 |
| Plasmopara halstedii GCA_900000015.1 | 218 | 0 | 2 | 14 |
| Pseudoperonospora humuli GCA_003991265.1 | 218 | 8 | 2 | 6 |
| Hyaloperonospora arabidopsidis GCA_001414525.1 | 215 | 2 | 5 | 12 |
| Plasmopara halstedii GCA_003640465.1 | 215 | 2 | 4 | 13 |
| Hyaloperonospora arabidopsidis GCA_001414265.1 | 213 | 2 | 7 | 12 |
| Peronospora cubensis AHJF01000001.1 | 213 | 3 | 2 | 16 |
| Phytopythium vexans AKYC02000001.1 | 213 | 0 | 3 | 18 |
| Phytophthora capsici ADVJ01000001.1 | 212 | 0 | 5 | 17 |
| Hyaloperonospora arabidopsidis GCA_000173235.2 | 209 | 7 | 6 | 12 |
| Sclerospora graminicola GCA_002933675.1 | 209 | 1 | 2 | 22 |
| Peronospora belbahrii * | 207 | 3 | 5 | 19 |
| Plasmopara halstedii GCA_003724065.1 | 203 | 1 | 2 | 28 |
| Saprolegnia parasitica GCA_000151545.2 | 191 | 13 | 1 | 29 |
| Albugo candida CAJG01000001.1 | 186 | 2 | 2 | 44 |
| Albugo candida LGTM01000001.1 | 186 | 2 | 3 | 43 |
| Albugo candida CAIW01000001.1 | 184 | 2 | 2 | 46 |
| Albugo candida CAIX01000001.1 | 183 | 5 | 3 | 43 |
| Albugo candida JZXB01000001.1 | 183 | 3 | 4 | 44 |
| Albugo candida LGTN01000001.1 | 179 | 1 | 4 | 50 |
| Albugo candida CAIV01000001.1 | 178 | 2 | 3 | 51 |
| Plasmopara halstedii GCA_003640505.1 | 165 | 18 | 5 | 46 |
| Sclerospora graminicola GCA_001887855.2 | 162 | 25 | 3 | 44 |
| Peronospora tabacina GCA_002099245.1 | 145 | 64 | 1 | 24 |
| Plasmopara viticola GCA_001974925.1 | 144 | 11 | 1 | 78 |
| Plasmopara viticola GCA_003123765.1 | 141 | 0 | 7 | 86 |
| Plasmopara obducens GCA_003640485.1 | 100 | 18 | 9 | 107 |
| Peronospora belbahrii GCA_002864105.1 | 94 | 126 | 4 | 10 |
| Pythium ultimum AKYB02000001.1 | 73 | 0 | 7 | 154 |
| Plasmopara obducens GCA_003640625.1 | 57 | 134 | 9 | 34 |
