## Supplementary material for "The genome of *Peronospora belbahrii* reveals high heterozygosity, a low number of canonical effectors and CT-rich promoters": All 19 supplementary files: Supplementary_Figure_5_gene_prediction.pptx

### Slide 1
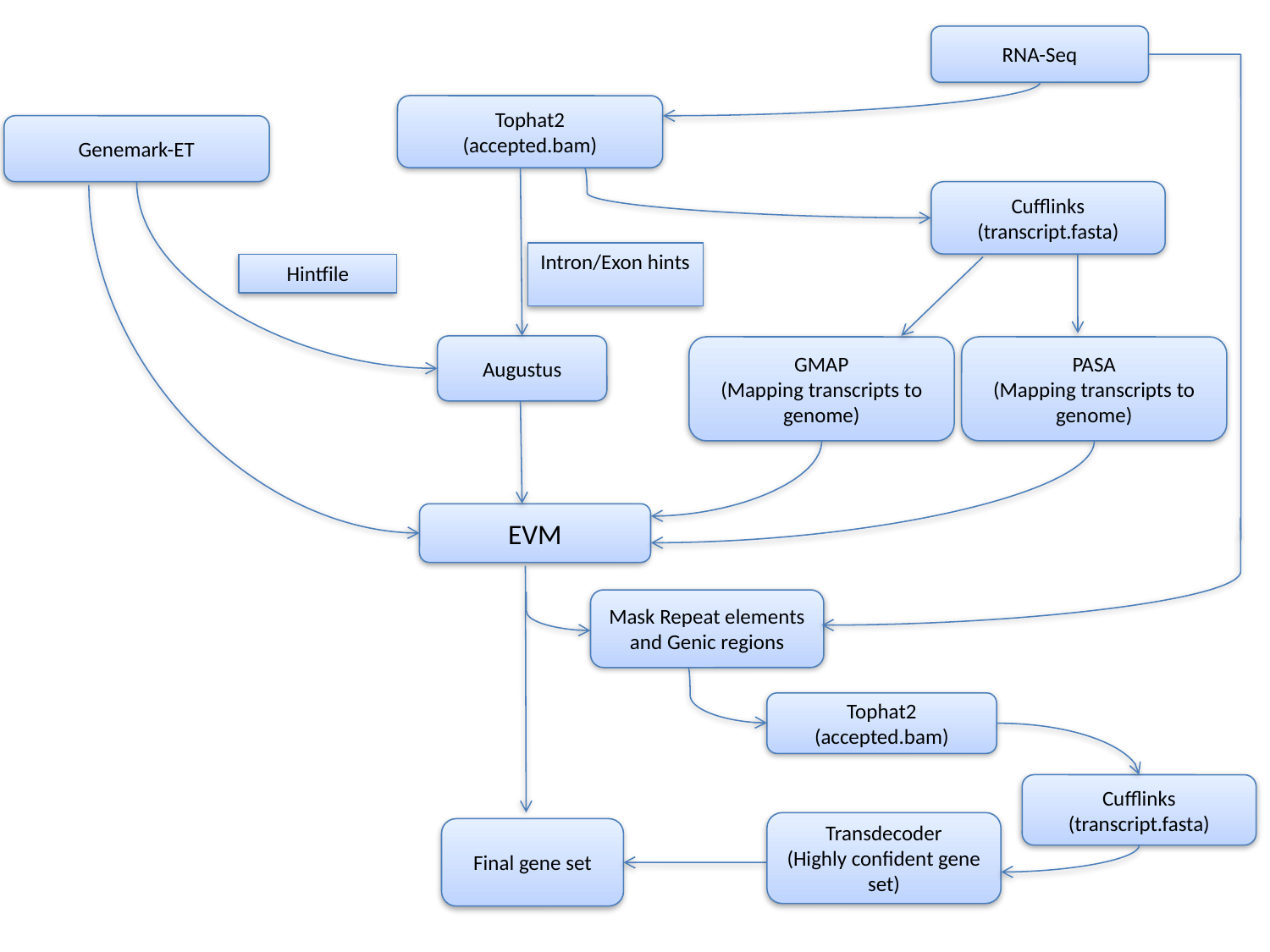

RNA-Seq
Tophat2
(accepted.bam)
Genemark-ET
Cufflinks
(transcript.fasta)
Intron/Exon hints
Hintfile
Augustus
GMAP
(Mapping transcripts to genome)
PASA
(Mapping transcripts to genome)
EVM
Mask Repeat elements and Genic regions
Tophat2
(accepted.bam)
Cufflinks
(transcript.fasta)
Transdecoder
(Highly confident gene set)
Final gene set
