## Supplementary material for "The genome of *Peronospora belbahrii* reveals high heterozygosity, a low number of canonical effectors and CT-rich promoters": All 19 supplementary files: Supplementary_Table_1_summary_Peronospora_genomes.docx

**Supplementary Table 1. Summary statistics for *Peronospora* genome sequence assemblies**.

| **Organism** | **Accession number** | **Reference** | **Scaffolds** | **Total length (b.p.)** | **Scaffold N_50_ (b.p.)** | **Scaffold L_50_** |
| --- | --- | --- | --- | --- | --- | --- |
| *P. belbahrii* | PRJEB20871 | This study | 2,780 | 35,394,047 | 248,464 | 46 |
| *P. belbahrii* | GCA_002864105.1 | Unpublished | 8,049 | 59,242,510 | 25,807 | 646 |
| *P. effusa* Pfs12 2209 | GCA_002245715.1 | (Feng et al. 2018) | 4,061 | 25,220,599 | 21,343 | 346 |
| *P. effusa* Pfs13 510c | GCA_002245725.1 | (Feng et al. 2018) | 4,387 | 23,992,579 | 18,134 | 378 |
| *P. effusa* Pfs14 4410 | GCA_002245735.1 | (Feng et al. 2018) | 4,020 | 24,880,192 | 22,141 | 327 |
| *P. effusa* R13 | GCA_003843895.1 | (Fletcher et al. 2018) | 784 | 32,162,939 | 72,179 | 204 |
| *P. effusa* R14 | GCA_003704535.1 | (Fletcher et al. 2018) | 880 | 30,813,317 | 61,367 | 161 |
| *P. tabacina* 968 S26 | GCA_002099245.1 | (Derevnina et al. 2015) | 4,015 | 63,065,868 | 78,591 | 918 |
