## Supplementary material for "The genome of *Peronospora belbahrii* reveals high heterozygosity, a low number of canonical effectors and CT-rich promoters": All 19 supplementary files: Supplementary_Table_2_summary_sequencing_reads.docx

Supplementary Table 2. Summary statistics for Illumina HiSeq 2000 sequencing data and quality control.

| **Sample** | **SRA accession number** | **Average depth of coverage** | **Number of read-pairs** | **Number of reads** | **Reads mapping to genome assembly** | |
| --- | --- | --- | --- | --- | --- | --- |
|  |  |  |  |  | **Number** | **Percentage** |
| Genomic paired-end (300 bp) | ERX1649693 | 111.98 x | 22,411,916 | 44,823,832 | 41,525,755 | 92.64% |
| Genomic paired-end  (800 bp) | ERX1649694 | 57.23 x | 11,488,769 | 22,977,538 | 21,217,402 | 92.34% |
| Genomic mate-pair (3 kbp) | ERX1649695 | 52.06 x | 22,936,656 | 45,873,312 | 20,727,967 | 45.19% |
| Genomic mate-pair  (8 kbp | ERX1649696 | 49.65 x | 14,031,175 | 28,062,350 | 19,858,590 | 70.77% |
| RNA-seq library | ERX1649697 | 9.39 x | 2,8528,694 | 57,057,388 | 14,981,720 | 26.26% |
