## Supplementary material for "The genome of *Peronospora belbahrii* reveals high heterozygosity, a low number of canonical effectors and CT-rich promoters": All 19 supplementary files: Supplementary_Table_3_genomes_for_heterozygosity_analyses.docx

Supplementary Table 3. Oomycete genomes used for heterozygosity estimation.

| Species | Accession numbers | Citation |
| --- | --- | --- |
| *Albugo candida* | SRR1811471 / CAIW01000001.1 | Links et al. 2011 |
| *Hyaloperonospora arabidopsidis* | ERR015782-ERR015789 / GCA_000173235.2 | Baxter et al. 2010 |
| *Peronospora tabacina* | SRR2146895 / Ptab_968-S26 | Derevnina et al. 2015 |
| *Phytophthora capsici* | SRR945695 / ADVJ01000001.1 | Lamour et al. 2012 |
| *Phytophthora infestans* | ERR248813 / GCA_000142945.1 | Haas et al. 2009 |
| *Phytophthora kernoviae* | SRR639379 GCA_000785735.2 | Studholme et al. 2016 |
| *Phytophthora lateralis* | SRR610738 / GCA_000318465.2 | Quinn et al. 2013 |
| *Phytophthora ramorum* | SRR639372 / GCA_000149735.1 | Tyler et a l. 2006 |
| *Phytophthora sojae* | SRR1046799 / GCA_000149755.2 | Tyler et al. 2006 |
| *Phytoythium vexans* | SRR235485 / AKYC02000001.1 | Adhikari et al. 2013 |
| *Plasmopara halstedii* | GCA_900000015.1 | Sharma et al. 2015 |
| *Pseudoperonospora cubensis* | SRR412826 / AHJF01000001.1 | Tian et al. 2011) |
| *Pythium ultimum* | SRR235484 / AKYB02000001.1 | Lévesque et al. 2010 |
| *Saprolegnia parasitica* | SRR058717 / GCA_000151545.2 | Jiang et al. 2013 |

Adhikari, B. N., Hamilton, J. P., Zerillo, M. M., Tisserat, N., Lévesque, C. A., and Buell, C. R. 2013. Comparative genomics reveals insight into virulence strategies of plant pathogenic oomycetes. ed. Boris Alexander Vinatzer. PLoS One. 8:e75072 Available at: http://dx.plos.org/10.1371/journal.pone.0075072 [Accessed October 22, 2013].

Baxter, L., Tripathy, S., Ishaque, N., Boot, N., Cabral, A., Kemen, E., et al. 2010. Signatures of Adaptation to Obligate Biotrophy in the Hyaloperonospora arabidopsidis Genome. Science (80-. ). 330:1549–1551. http://www.sciencemag.org/content/330/6010/1549.short.

Derevnina, L., Chin-Wo-Reyes, S., Martin, F., Wood, K., Froenicke, L., Spring, O., et al. 2015. Genome Sequence and Architecture of the Tobacco Downy Mildew Pathogen, Peronospora tabacina. Mol. Plant-Microbe Interact. 28:150721061836007 Available at: http://apsjournals.apsnet.org/doi/10.1094/MPMI-05-15-0112-R.

Feng, C., Lamour, K. H., Bluhm, B. H., Sharma, S., Shrestha, S., Dhillon, B. D. S., et al. 2018. Genome Sequences of Three Races of *Peronospora effusa* : A Resource for Studying the Evolution of the Spinach Downy Mildew Pathogen. Mol. Plant-Microbe Interact. 31:1230–1231 Available at: https://apsjournals.apsnet.org/doi/10.1094/MPMI-04-18-0085-A.

Fletcher, K., Klosterman, S. J., Derevnina, L., Martin, F., Bertier, L. D., Koike, S., et al. 2018. Comparative genomics of downy mildews reveals potential adaptations to biotrophy. BMC Genomics. 19:851 Available at: https://bmcgenomics.biomedcentral.com/articles/10.1186/s12864-018-5214-8.

Haas, B. J. B. J., Kamoun, S., Zody, M. C., Jiang, R. H. Y. R. H. Y., Handsaker, R. E. R. E., Cano, L. M. L. M., et al. 2009. Genome sequence and analysis of the Irish potato famine pathogen Phytophthora infestans. Nature. 461:393–398 Available at: http://www.nature.com/nature/journal/v461/n7262/full/nature08358.html.

Jiang, R. H. Y., de Bruijn, I., Haas, B. J., Belmonte, R., Löbach, L., Christie, J., et al. 2013. Distinctive Expansion of Potential Virulence Genes in the Genome of the Oomycete Fish Pathogen Saprolegnia parasitica ed. John M. McDowell. PLoS Genet. 9:e1003272 Available at: http://dx.plos.org/10.1371/journal.pgen.1003272 [Accessed June 14, 2013].

Lamour, K. H., Mudge, J., Gobena, D., Hurtado-Gonzales, O. P., Schmutz, J., Kuo, A., et al. 2012. Genome sequencing and mapping reveal loss of heterozygosity as a mechanism for rapid adaptation in the vegetable pathogen Phytophthora capsici. Mol. Plant. Microbe. Interact. 25:1350–60 Available at: http://www.ncbi.nlm.nih.gov/pubmed/22712506 [Accessed January 20, 2013].

Lévesque, C. A., Brouwer, H., Cano, L., Hamilton, J. P., Holt, C., Huitema, E., et al. 2010. Genome sequence of the necrotrophic plant pathogen Pythium ultimum reveals original pathogenicity mechanisms and effector repertoire. Genome Biol. 11:R73 Available at: http://genomebiology.com/2010/11/7/R73 [Accessed November 3, 2013].

Li, H., and Durbin, R. 2009. Fast and accurate short read alignment with Burrows-Wheeler transform. Bioinformatics. 25:1754–60 Available at: http://www.pubmedcentral.nih.gov/articlerender.fcgi?artid=2705234&tool=pmcentrez&rendertype=abstract [Accessed November 1, 2012].

Links, M. G., Holub, E., Jiang, R. H. Y., Sharpe, A. G., Hegedus, D., Beynon, E., et al. 2011. De novo sequence assembly of Albugo candida reveals a small genome relative to other biotrophic oomycetes. BMC Genomics. 12:503 Available at: http://www.pubmedcentral.nih.gov/articlerender.fcgi?artid=3206522&tool=pmcentrez&rendertype=abstract [Accessed October 26, 2012].

Quinn, L., O’Neill, P. a, Harrison, J., Paskiewicz, K. H., McCracken, A. R., Cooke, L. R., et al. 2013. Genome-wide sequencing of Phytophthora lateralis reveals genetic variation among isolates from Lawson cypress ( Chamaecyparis lawsoniana ) in Northern Ireland. FEMS Microbiol. Lett. 344:179–185 Available at: http://www.ncbi.nlm.nih.gov/pubmed/23678994 [Accessed June 8, 2013].

Savory, E. a, Zou, C., Adhikari, B. N., Hamilton, J. P., Buell, C. R., Shiu, S.-H., et al. 2012. Alternative splicing of a multi-drug transporter from Pseudoperonospora cubensis generates an RXLR effector protein that elicits a rapid cell death. PLoS One. 7:e34701 Available at: http://www.pubmedcentral.nih.gov/articlerender.fcgi?artid=3320632&tool=pmcentrez&rendertype=abstract.

Sharma, R., Xia, X., Cano, L. M., Evangelisti, E., Kemen, E., Judelson, H., et al. 2015. Genome analyses of the sunflower pathogen Plasmopara halstedii provide insights into effector evolution in downy mildews and Phytophthora. BMC Genomics. 16:741 Available at: http://www.biomedcentral.com/1471-2164/16/741.

Studholme, D. J., McDougal, R. L., Sambles, C., Hansen, E., Hardy, G., Grant, M., et al. 2016. Genome sequences of six Phytophthora species associated with forests in New Zealand. Genomics Data. 7:54–56 Available at: http://linkinghub.elsevier.com/retrieve/pii/S2213596015300854.

Thorvaldsdottir, H., Robinson, J. T., and Mesirov, J. P. 2013. Integrative Genomics Viewer (IGV): high-performance genomics data visualization and exploration. Brief. Bioinform. 14:178–192 Available at: http://bib.oxfordjournals.org/cgi/doi/10.1093/bib/bbs017.

Tian, M., Win, J., Savory, E., Burkhardt, A., Held, M., Brandizzi, F., et al. 2011. 454 Genome sequencing of Pseudoperonospora cubensis reveals effector proteins with a QXLR translocation motif. Mol. Plant. Microbe. Interact. 24:543–553.

Tyler, B. M., Tripathy, S., Zhang, X., Dehal, P., Jiang, R. H. Y., Aerts, A., et al. 2006. Phytophthora genome sequences uncover evolutionary origins and mechanisms of pathogenesis. Science. 313:1261–1266 Available at: http://www.sciencemag.org/content/313/5791/1261.short.
