## Supplementary material for "The genome of *Peronospora belbahrii* reveals high heterozygosity, a low number of canonical effectors and CT-rich promoters": All 19 supplementary files: Supplementary_Table_4_metabolic_reactions_supported.docx

Supplementary Table 4. Supported metabolic reactions.

| **KEGG pathway ID** | **KEGG BRITE category** | **KEGG pathway** | **total reactions** | **Peronospora belbahrii** | **Plasmopara halstedii** | **Phytophthora infestans** | **Phytophthora capsici** | **Phytophthora sojae** | **Phytophthora ramorum** | **Phytopythium vexans** | **Hyaloperonospora arabidopsidis** | **Pythium ultimum** | **Albugo laibachii** | **Saprolegnia parasitica** | **variance** | **mean** | **var/mean** |
| --- | --- | --- | --- | --- | --- | --- | --- | --- | --- | --- | --- | --- | --- | --- | --- | --- | --- |
| **100** | **Lipid metabolism** | **Steroid biosynthesis** | **54** | **1** | **1** | **10** | **9** | **10** | **10** | **10** | **1** | **6** | **2** | **33** | **82,67272727** | **8,454545455** | **9,778494624** |
| **1100** | **Global and overview maps** | **Metabolic pathways** | **2411** | **650** | **691** | **751** | **760** | **757** | **705** | **749** | **643** | **544** | **596** | **762** | **5593,254545** | **691,6363636** | **8,086987382** |
| **1110** | **Global and overview maps** | **Biosynthesis of secondary metabolites** | **1201** | **244** | **258** | **268** | **276** | **273** | **251** | **275** | **232** | **181** | **242** | **287** | **864,8545455** | **253,3636364** | **3,413491209** |
| **120** | **Lipid metabolism** | **Primary bile acid biosynthesis** | **53** | **10** | **8** | **8** | **8** | **8** | **4** | **8** | **10** | **6** | **4** | **21** | **20,85454545** | **8,636363636** | **2,414736842** |
| **940** | **Biosynthesis of other secondary metabolites** | **Phenylpropanoid biosynthesis** | **87** | **5** | **16** | **20** | **20** | **22** | **20** | **20** | **9** | **11** | **12** | **22** | **34,69090909** | **16,09090909** | **2,155932203** |
| **140** | **Lipid metabolism** | **Steroid hormone biosynthesis** | **139** | **15** | **11** | **13** | **15** | **15** | **15** | **13** | **9** | **2** | **10** | **21** | **22,85454545** | **12,63636364** | **1,808633094** |
| **40** | **Carbohydrate metabolism** | **Pentose and glucuronate interconversions** | **73** | **8** | **8** | **15** | **15** | **15** | **14** | **12** | **7** | **7** | **5** | **8** | **14,85454545** | **10,36363636** | **1,433333333** |
| **130** | **Metabolism of cofactors and vitamins** | **Ubiquinone and other terpenoid-quinone biosynthesis** | **89** | **19** | **20** | **20** | **20** | **20** | **10** | **17** | **11** | **7** | **19** | **18** | **22,67272727** | **16,45454545** | **1,377900552** |
| **350** | **Amino acid metabolism** | **Tyrosine metabolism** | **105** | **23** | **18** | **26** | **29** | **30** | **31** | **31** | **20** | **26** | **14** | **22** | **32,07272727** | **24,54545455** | **1,306666667** |
| **240** | **Nucleotide metabolism** | **Pyrimidine metabolism** | **119** | **26** | **26** | **35** | **35** | **35** | **25** | **29** | **21** | **22** | **18** | **29** | **34,65454545** | **27,36363636** | **1,266445183** |
| **230** | **Nucleotide metabolism** | **Purine metabolism** | **152** | **59** | **68** | **76** | **76** | **77** | **69** | **71** | **69** | **50** | **55** | **64** | **79,21818182** | **66,72727273** | **1,18719346** |
| **830** | **Metabolism of cofactors and vitamins** | **Retinol metabolism** | **31** | **6** | **7** | **8** | **11** | **11** | **10** | **8** | **11** | **9** | **1** | **11** | **9,272727273** | **8,454545455** | **1,096774194** |
| **626** | **Xenobiotics biodegradation and metabolism** | **Naphthalene degradation** | **59** | **6** | **2** | **6** | **6** | **6** | **6** | **6** | **2** | **2** | **2** | **2** | **4,363636364** | **4,181818182** | **1,043478261** |
| **950** | **Biosynthesis of other secondary metabolites** | **Isoquinoline alkaloid biosynthesis** | **100** | **1** | **1** | **1** | **4** | **4** | **4** | **4** | **1** | **1** | **1** | **3** | **2,218181818** | **2,272727273** | **0,976** |
| **330** | **Amino acid metabolism** | **Arginine and proline metabolism** | **107** | **19** | **29** | **31** | **32** | **30** | **29** | **27** | **25** | **20** | **23** | **31** | **20,69090909** | **26,90909091** | **0,768918919** |
| **480** | **Metabolism of other amino acids** | **Glutathione metabolism** | **53** | **20** | **23** | **22** | **23** | **23** | **23** | **23** | **23** | **15** | **21** | **32** | **15,67272727** | **22,54545455** | **0,69516129** |
| **983** | **Xenobiotics biodegradation and metabolism** | **Drug metabolism - other enzymes** | **38** | **8** | **8** | **8** | **12** | **12** | **11** | **11** | **7** | **8** | **4** | **8** | **5,963636364** | **8,818181818** | **0,67628866** |
| **750** | **Metabolism of cofactors and vitamins** | **Vitamin B6 metabolism** | **39** | **5** | **9** | **9** | **6** | **9** | **3** | **8** | **5** | **6** | **5** | **7** | **4,072727273** | **6,545454545** | **0,622222222** |
| **270** | **Amino acid metabolism** | **Cysteine and methionine metabolism** | **88** | **36** | **37** | **37** | **34** | **37** | **38** | **36** | **36** | **23** | **29** | **32** | **20,49090909** | **34,09090909** | **0,601066667** |
| **960** | **Biosynthesis of other secondary metabolites** | **Tropane, piperidine and pyridine alkaloid biosynthesis** | **68** | **1** | **1** | **1** | **3** | **3** | **3** | **3** | **1** | **1** | **1** | **1** | **1,018181818** | **1,727272727** | **0,589473684** |
| **260** | **Amino acid metabolism** | **Glycine, serine and threonine metabolism** | **71** | **35** | **33** | **37** | **39** | **35** | **34** | **38** | **32** | **23** | **31** | **37** | **19,6** | **34** | **0,576470588** |
| **590** | **Lipid metabolism** | **Arachidonic acid metabolism** | **62** | **6** | **6** | **10** | **10** | **10** | **10** | **10** | **6** | **6** | **5** | **9** | **4,6** | **8** | **0,575** |
| **643** | **Xenobiotics biodegradation and metabolism** | **Styrene degradation** | **23** | **5** | **4** | **6** | **6** | **6** | **4** | **5** | **6** | **7** | **1** | **6** | **2,690909091** | **5,090909091** | **0,528571429** |
| **591** | **Lipid metabolism** | **Linoleic acid metabolism** | **16** | **3** | **1** | **1** | **3** | **3** | **3** | **1** | **3** | **1** | **1** | **3** | **1,090909091** | **2,090909091** | **0,52173913** |
| **562** | **Carbohydrate metabolism** | **Inositol phosphate metabolism** | **57** | **27** | **28** | **25** | **27** | **25** | **21** | **25** | **25** | **16** | **23** | **27** | **11,87272727** | **24,45454545** | **0,485501859** |
| **625** | **Xenobiotics biodegradation and metabolism** | **Chloroalkane and chloroalkene degradation** | **45** | **5** | **5** | **6** | **9** | **6** | **6** | **6** | **5** | **5** | **2** | **6** | **2,672727273** | **5,545454545** | **0,481967213** |
| **430** | **Metabolism of other amino acids** | **Taurine and hypotaurine metabolism** | **23** | **4** | **7** | **8** | **8** | **8** | **8** | **6** | **5** | **3** | **6** | **6** | **3,018181818** | **6,272727273** | **0,48115942** |
| **400** | **Amino acid metabolism** | **Phenylalanine, tyrosine and tryptophan biosynthesis** | **49** | **20** | **21** | **21** | **21** | **21** | **14** | **24** | **21** | **14** | **21** | **21** | **9,490909091** | **19,90909091** | **0,476712329** |
| **340** | **Amino acid metabolism** | **Histidine metabolism** | **53** | **10** | **9** | **12** | **13** | **13** | **10** | **13** | **7** | **8** | **9** | **13** | **5,054545455** | **10,63636364** | **0,475213675** |
| **740** | **Metabolism of cofactors and vitamins** | **Riboflavin metabolism** | **26** | **6** | **8** | **6** | **6** | **5** | **6** | **6** | **5** | **2** | **4** | **4** | **2,418181818** | **5,272727273** | **0,45862069** |
| **900** | **Metabolism of terpenoids and polyketides** | **Terpenoid backbone biosynthesis** | **56** | **15** | **16** | **16** | **16** | **14** | **13** | **16** | **15** | **8** | **15** | **18** | **6,618181818** | **14,72727273** | **0,449382716** |
| **52** | **Carbohydrate metabolism** | **Galactose metabolism** | **57** | **11** | **11** | **15** | **15** | **15** | **15** | **13** | **11** | **12** | **11** | **8** | **5,472727273** | **12,45454545** | **0,439416058** |
| **860** | **Metabolism of cofactors and vitamins** | **Porphyrin and chlorophyll metabolism** | **141** | **13** | **12** | **16** | **16** | **16** | **16** | **15** | **12** | **9** | **11** | **15** | **6,018181818** | **13,72727273** | **0,438410596** |
| **790** | **Metabolism of cofactors and vitamins** | **Folate biosynthesis** | **40** | **23** | **25** | **23** | **23** | **25** | **23** | **22** | **23** | **15** | **18** | **25** | **9,618181818** | **22,27272727** | **0,431836735** |
| **510** | **Glycan biosynthesis and metabolism** | **N-Glycan biosynthesis** | **35** | **20** | **23** | **22** | **20** | **23** | **14** | **22** | **21** | **16** | **22** | **22** | **8,472727273** | **20,45454545** | **0,414222222** |
| **1210** | **Global and overview maps** | **2-Oxocarboxylic acid metabolism** | **131** | **36** | **35** | **36** | **34** | **35** | **31** | **36** | **32** | **24** | **33** | **37** | **13,47272727** | **33,54545455** | **0,401626016** |
| **360** | **Amino acid metabolism** | **Phenylalanine metabolism** | **85** | **8** | **9** | **10** | **12** | **12** | **12** | **11** | **9** | **8** | **6** | **10** | **3,818181818** | **9,727272727** | **0,392523364** |
| **290** | **Amino acid metabolism** | **Valine, leucine and isoleucine biosynthesis** | **25** | **15** | **15** | **15** | **15** | **15** | **15** | **15** | **11** | **8** | **15** | **15** | **5,4** | **14** | **0,385714286** |
| **910** | **Energy metabolism** | **Nitrogen metabolism** | **42** | **9** | **8** | **13** | **13** | **13** | **11** | **10** | **9** | **13** | **9** | **9** | **4,054545455** | **10,63636364** | **0,381196581** |
| **73** | **Lipid metabolism** | **Cutin, suberine and wax biosynthesis** | **26** | **5** | **5** | **5** | **4** | **5** | **4** | **6** | **4** | **4** | **4** | **1** | **1,618181818** | **4,272727273** | **0,378723404** |
| **627** | **Xenobiotics biodegradation and metabolism** | **Aminobenzoate degradation** | **106** | **7** | **7** | **7** | **7** | **7** | **6** | **7** | **7** | **8** | **2** | **7** | **2,472727273** | **6,545454545** | **0,377777778** |
| **760** | **Metabolism of cofactors and vitamins** | **Nicotinate and nicotinamide metabolism** | **68** | **8** | **13** | **13** | **13** | **13** | **13** | **12** | **12** | **8** | **9** | **12** | **4,272727273** | **11,45454545** | **0,373015873** |
| **930** | **Xenobiotics biodegradation and metabolism** | **Caprolactam degradation** | **22** | **3** | **3** | **4** | **4** | **4** | **4** | **3** | **2** | **1** | **3** | **5** | **1,218181818** | **3,272727273** | **0,372222222** |
| **1200** | **Global and overview maps** | **Carbon metabolism** | **194** | **82** | **81** | **86** | **86** | **84** | **81** | **86** | **81** | **68** | **75** | **83** | **29,36363636** | **81,18181818** | **0,361702128** |
| **561** | **Lipid metabolism** | **Glycerolipid metabolism** | **42** | **13** | **13** | **15** | **14** | **15** | **14** | **14** | **12** | **9** | **9** | **13** | **4,363636364** | **12,81818182** | **0,340425532** |
| **520** | **Carbohydrate metabolism** | **Amino sugar and nucleotide sugar metabolism** | **138** | **16** | **16** | **20** | **21** | **20** | **20** | **21** | **16** | **15** | **16** | **19** | **5,563636364** | **18,18181818** | **0,306** |
| **51** | **Carbohydrate metabolism** | **Fructose and mannose metabolism** | **79** | **12** | **11** | **15** | **16** | **15** | **16** | **14** | **12** | **11** | **11** | **14** | **4,054545455** | **13,36363636** | **0,303401361** |
| **460** | **Metabolism of other amino acids** | **Cyanoamino acid metabolism** | **51** | **7** | **9** | **9** | **9** | **10** | **10** | **14** | **9** | **10** | **9** | **9** | **2,872727273** | **9,545454545** | **0,300952381** |
| **563** | **Glycan biosynthesis and metabolism** | **Glycosylphosphatidylinositol(GPI)-anchor biosynthesis** | **11** | **9** | **8** | **8** | **5** | **8** | **5** | **9** | **9** | **9** | **8** | **8** | **2,163636364** | **7,818181818** | **0,276744186** |
| **720** | **Energy metabolism** | **Carbon fixation pathways in prokaryotes** | **55** | **21** | **20** | **23** | **23** | **22** | **19** | **23** | **20** | **19** | **15** | **21** | **5,672727273** | **20,54545455** | **0,276106195** |
| **450** | **Metabolism of other amino acids** | **Selenocompound metabolism** | **24** | **13** | **12** | **15** | **13** | **14** | **13** | **14** | **14** | **10** | **9** | **14** | **3,363636364** | **12,81818182** | **0,262411348** |
| **730** | **Metabolism of cofactors and vitamins** | **Thiamine metabolism** | **28** | **3** | **3** | **3** | **3** | **3** | **3** | **3** | **3** | **3** | **3** | **6** | **0,818181818** | **3,272727273** | **0,25** |
| **380** | **Amino acid metabolism** | **Tryptophan metabolism** | **95** | **13** | **15** | **15** | **18** | **18** | **17** | **18** | **15** | **15** | **13** | **18** | **3,890909091** | **15,90909091** | **0,244571429** |
| **600** | **Lipid metabolism** | **Sphingolipid metabolism** | **40** | **11** | **14** | **12** | **14** | **14** | **12** | **13** | **10** | **12** | **9** | **11** | **2,8** | **12** | **0,233333333** |
| **62** | **Lipid metabolism** | **Fatty acid elongation** | **35** | **33** | **32** | **33** | **33** | **33** | **33** | **33** | **32** | **24** | **33** | **33** | **7,2** | **32** | **0,225** |
| **500** | **Carbohydrate metabolism** | **Starch and sucrose metabolism** | **84** | **16** | **18** | **18** | **18** | **18** | **18** | **15** | **18** | **13** | **15** | **14** | **3,672727273** | **16,45454545** | **0,22320442** |
| **780** | **Metabolism of cofactors and vitamins** | **Biotin metabolism** | **24** | **9** | **9** | **9** | **9** | **9** | **11** | **11** | **7** | **7** | **7** | **9** | **1,963636364** | **8,818181818** | **0,222680412** |
| **680** | **Energy metabolism** | **Methane metabolism** | **89** | **18** | **18** | **19** | **19** | **18** | **18** | **19** | **18** | **12** | **17** | **18** | **3,854545455** | **17,63636364** | **0,218556701** |
| **30** | **Carbohydrate metabolism** | **Pentose phosphate pathway** | **54** | **19** | **20** | **22** | **22** | **22** | **22** | **21** | **20** | **15** | **20** | **20** | **4,218181818** | **20,27272727** | **0,208071749** |
| **53** | **Carbohydrate metabolism** | **Ascorbate and aldarate metabolism** | **59** | **7** | **7** | **9** | **9** | **9** | **9** | **8** | **6** | **7** | **6** | **9** | **1,563636364** | **7,818181818** | **0,2** |
| **521** | **Biosynthesis of other secondary metabolites** | **Streptomycin biosynthesis** | **24** | **6** | **6** | **6** | **6** | **5** | **6** | **6** | **5** | **3** | **4** | **5** | **1,018181818** | **5,272727273** | **0,193103448** |
| **300** | **Amino acid metabolism** | **Lysine biosynthesis** | **39** | **10** | **10** | **10** | **10** | **10** | **9** | **10** | **9** | **6** | **11** | **11** | **1,854545455** | **9,636363636** | **0,19245283** |
| **630** | **Carbohydrate metabolism** | **Glyoxylate and dicarboxylate metabolism** | **81** | **22** | **21** | **24** | **24** | **24** | **21** | **25** | **22** | **18** | **21** | **24** | **4,254545455** | **22,36363636** | **0,190243902** |
| **410** | **Metabolism of other amino acids** | **beta-Alanine metabolism** | **37** | **12** | **12** | **12** | **13** | **13** | **13** | **13** | **12** | **8** | **11** | **12** | **2,090909091** | **11,90909091** | **0,175572519** |
| **261** | **Biosynthesis of other secondary metabolites** | **Monobactam biosynthesis** | **28** | **3** | **3** | **3** | **3** | **3** | **3** | **3** | **2** | **1** | **3** | **2** | **0,454545455** | **2,636363636** | **0,172413793** |
| **1212** | **Global and overview maps** | **Fatty acid metabolism** | **98** | **84** | **83** | **84** | **84** | **84** | **84** | **84** | **83** | **72** | **83** | **84** | **12,65454545** | **82,63636364** | **0,153135314** |
| **920** | **Energy metabolism** | **Sulfur metabolism** | **47** | **6** | **7** | **9** | **8** | **8** | **8** | **9** | **7** | **7** | **6** | **7** | **1,072727273** | **7,454545455** | **0,143902439** |
| **592** | **Lipid metabolism** | **alpha-Linolenic acid metabolism** | **31** | **7** | **8** | **8** | **9** | **8** | **8** | **8** | **8** | **5** | **7** | **8** | **1,054545455** | **7,636363636** | **0,138095238** |
| **640** | **Carbohydrate metabolism** | **Propanoate metabolism** | **66** | **19** | **19** | **21** | **21** | **21** | **22** | **21** | **21** | **17** | **18** | **22** | **2,763636364** | **20,18181818** | **0,136936937** |
| **250** | **Amino acid metabolism** | **Alanine, aspartate and glutamate metabolism** | **47** | **26** | **30** | **32** | **32** | **32** | **31** | **32** | **28** | **29** | **29** | **29** | **4** | **30** | **0,133333333** |
| **310** | **Amino acid metabolism** | **Lysine degradation** | **63** | **15** | **16** | **16** | **17** | **17** | **17** | **18** | **17** | **15** | **16** | **20** | **2,018181818** | **16,72727273** | **0,120652174** |
| **513** | **Glycan biosynthesis and metabolism** | **Various types of N-glycan biosynthesis** | **25** | **12** | **12** | **12** | **10** | **12** | **9** | **12** | **10** | **12** | **11** | **12** | **1,218181818** | **11,27272727** | **0,108064516** |
| **770** | **Metabolism of cofactors and vitamins** | **Pantothenate and CoA biosynthesis** | **32** | **14** | **15** | **14** | **14** | **13** | **14** | **16** | **12** | **12** | **14** | **15** | **1,490909091** | **13,90909091** | **0,107189542** |
| **10** | **Carbohydrate metabolism** | **Glycolysis / Gluconeogenesis** | **54** | **28** | **29** | **30** | **30** | **30** | **29** | **30** | **28** | **24** | **29** | **29** | **3,018181818** | **28,72727273** | **0,105063291** |
| **564** | **Lipid metabolism** | **Glycerophospholipid metabolism** | **87** | **38** | **39** | **41** | **39** | **40** | **40** | **40** | **36** | **35** | **36** | **38** | **3,854545455** | **38,36363636** | **0,100473934** |
| **1040** | **Lipid metabolism** | **Biosynthesis of unsaturated fatty acids** | **41** | **16** | **15** | **16** | **16** | **16** | **16** | **16** | **15** | **12** | **16** | **16** | **1,472727273** | **15,45454545** | **0,095294118** |
| **440** | **Metabolism of other amino acids** | **Phosphonate and phosphinate metabolism** | **52** | **3** | **3** | **3** | **3** | **2** | **3** | **3** | **3** | **3** | **4** | **4** | **0,290909091** | **3,090909091** | **0,094117647** |
| **280** | **Amino acid metabolism** | **Valine, leucine and isoleucine degradation** | **50** | **34** | **34** | **37** | **37** | **37** | **35** | **36** | **34** | **33** | **32** | **37** | **3,290909091** | **35,09090909** | **0,093782383** |
| **970** | **Translation** | **Aminoacyl-tRNA biosynthesis** | **32** | **25** | **24** | **26** | **26** | **24** | **24** | **25** | **25** | **21** | **23** | **26** | **2,272727273** | **24,45454545** | **0,092936803** |
| **670** | **Metabolism of cofactors and vitamins** | **One carbon pool by folate** | **33** | **20** | **20** | **21** | **21** | **19** | **21** | **20** | **20** | **17** | **18** | **21** | **1,763636364** | **19,81818182** | **0,088990826** |
| **220** | **Amino acid metabolism** | **Arginine biosynthesis** | **31** | **19** | **17** | **18** | **17** | **18** | **18** | **19** | **19** | **15** | **19** | **19** | **1,6** | **18** | **0,088888889** |
| **20** | **Carbohydrate metabolism** | **Citrate cycle (TCA cycle)** | **28** | **20** | **20** | **20** | **20** | **20** | **17** | **20** | **20** | **19** | **17** | **20** | **1,454545455** | **19,36363636** | **0,075117371** |
| **980** | **Xenobiotics biodegradation and metabolism** | **Metabolism of xenobiotics by cytochrome P450** | **121** | **3** | **3** | **3** | **3** | **3** | **3** | **2** | **3** | **2** | **3** | **3** | **0,163636364** | **2,818181818** | **0,058064516** |
| **982** | **Xenobiotics biodegradation and metabolism** | **Drug metabolism - cytochrome P450** | **84** | **6** | **7** | **7** | **8** | **7** | **7** | **8** | **7** | **7** | **6** | **7** | **0,4** | **7** | **0,057142857** |
| **620** | **Carbohydrate metabolism** | **Pyruvate metabolism** | **71** | **26** | **26** | **27** | **27** | **27** | **25** | **27** | **26** | **23** | **26** | **27** | **1,490909091** | **26,09090909** | **0,057142857** |
| **565** | **Lipid metabolism** | **Ether lipid metabolism** | **32** | **8** | **8** | **8** | **9** | **9** | **9** | **9** | **8** | **8** | **7** | **8** | **0,418181818** | **8,272727273** | **0,050549451** |
| **660** | **Carbohydrate metabolism** | **C5-Branched dibasic acid metabolism** | **34** | **2** | **2** | **2** | **2** | **2** | **2** | **2** | **2** | **1** | **2** | **2** | **0,090909091** | **1,909090909** | **0,047619048** |
| **281** | **Metabolism of terpenoids and polyketides** | **Geraniol degradation** | **23** | **2** | **2** | **2** | **2** | **2** | **2** | **2** | **2** | **1** | **2** | **2** | **0,090909091** | **1,909090909** | **0,047619048** |
| **650** | **Carbohydrate metabolism** | **Butanoate metabolism** | **63** | **13** | **13** | **14** | **14** | **14** | **13** | **13** | **13** | **12** | **13** | **14** | **0,418181818** | **13,27272727** | **0,031506849** |
| **332** | **Biosynthesis of other secondary metabolites** | **Carbapenem biosynthesis** | **29** | **3** | **3** | **3** | **3** | **3** | **3** | **3** | **3** | **2** | **3** | **3** | **0,090909091** | **2,909090909** | **0,03125** |
| **362** | **Xenobiotics biodegradation and metabolism** | **Benzoate degradation** | **87** | **2** | **3** | **3** | **3** | **3** | **3** | **3** | **3** | **3** | **3** | **3** | **0,090909091** | **2,909090909** | **0,03125** |
| **471** | **Metabolism of other amino acids** | **D-Glutamine and D-glutamate metabolism** | **11** | **3** | **3** | **3** | **3** | **3** | **3** | **3** | **3** | **3** | **3** | **4** | **0,090909091** | **3,090909091** | **0,029411765** |
| **72** | **Lipid metabolism** | **Synthesis and degradation of ketone bodies** | **6** | **4** | **4** | **4** | **4** | **4** | **4** | **3** | **4** | **4** | **4** | **4** | **0,090909091** | **3,909090909** | **0,023255814** |
| **710** | **Energy metabolism** | **Carbon fixation in photosynthetic organisms** | **26** | **18** | **18** | **19** | **19** | **19** | **18** | **19** | **18** | **17** | **18** | **18** | **0,418181818** | **18,27272727** | **0,022885572** |
| **71** | **Lipid metabolism** | **Fatty acid degradation** | **47** | **33** | **34** | **34** | **34** | **34** | **34** | **34** | **34** | **32** | **33** | **34** | **0,454545455** | **33,63636364** | **0,013513514** |
| **61** | **Lipid metabolism** | **Fatty acid biosynthesis** | **57** | **35** | **35** | **35** | **35** | **35** | **35** | **35** | **35** | **33** | **34** | **35** | **0,418181818** | **34,72727273** | **0,012041885** |
| **254** | **Biosynthesis of other secondary metabolites** | **Aflatoxin biosynthesis** | **29** | **1** | **1** | **1** | **1** | **1** | **1** | **1** | **1** | **1** | **1** | **1** | **0** | **1** | **0** |
| **253** | **Metabolism of terpenoids and polyketides** | **Tetracycline biosynthesis** | **22** | **1** | **1** | **1** | **1** | **1** | **1** | **1** | **1** | **1** | **1** | **1** | **0** | **1** | **0** |
| **524** | **Biosynthesis of other secondary metabolites** | **Butirosin and neomycin biosynthesis** | **30** | **1** | **1** | **1** | **1** | **1** | **1** | **1** | **1** | **1** | **1** | **1** | **0** | **1** | **0** |
| **402** | **Biosynthesis of other secondary metabolites** | **Benzoxazinoid biosynthesis** | **8** | **1** | **1** | **1** | **1** | **1** | **1** | **1** | **1** | **1** | **1** | **1** | **0** | **1** | **0** |
| **966** | **Biosynthesis of other secondary metabolites** | **Glucosinolate biosynthesis** | **59** | **3** | **3** | **3** | **3** | **3** | **3** | **3** | **3** | **3** | **3** | **3** | **0** | **3** | **0** |
| **514** | **Glycan biosynthesis and metabolism** | **Other types of O-glycan biosynthesis** | **18** | **1** | **1** | **1** | **1** | **1** | **1** | **1** | **1** | **1** | **1** | **1** | **0** | **1** | **0** |
| **401** | **Biosynthesis of other secondary metabolites** | **Novobiocin biosynthesis** | **31** | **1** | **1** | **1** | **1** | **1** | **1** | **1** | **1** | **1** | **1** | **1** | **0** | **1** | **0** |
| **1051** | **Metabolism of terpenoids and polyketides** | **Biosynthesis of ansamycins** | **30** | **1** | **1** | **1** | **1** | **1** | **1** | **1** | **1** | **1** | **1** | **1** | **0** | **1** | **0** |
