## Supplementary figures and images for "The genome of *Peronospora belbahrii* reveals high heterozygosity, a low number of canonical effectors and CT-rich promoters"

### Supplementary_Figure_1_mitogenome_circle.pptx

#### Slide 1
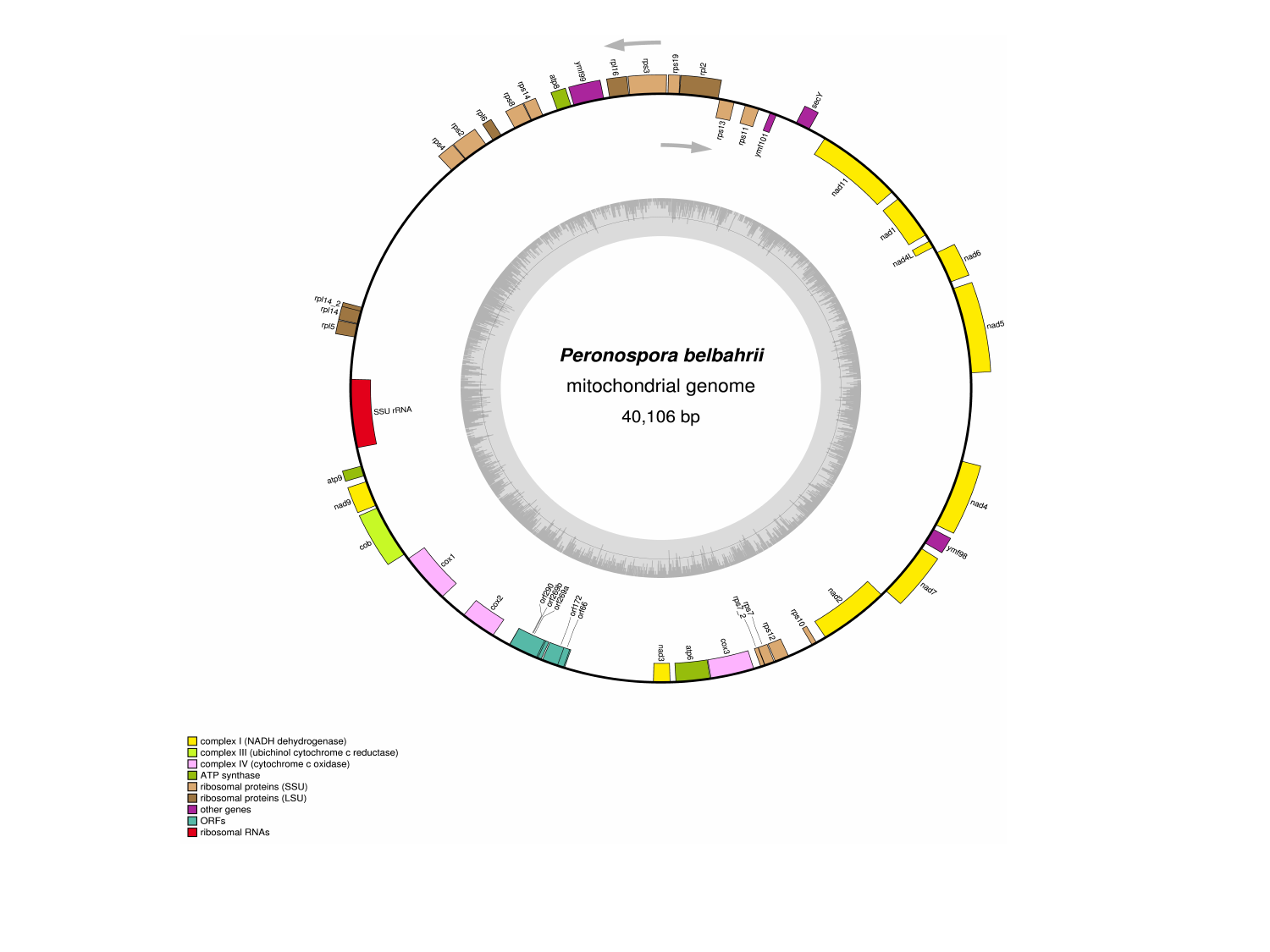

### Supplementary_Figure_3_promotor_features.pptx

#### Slide 1
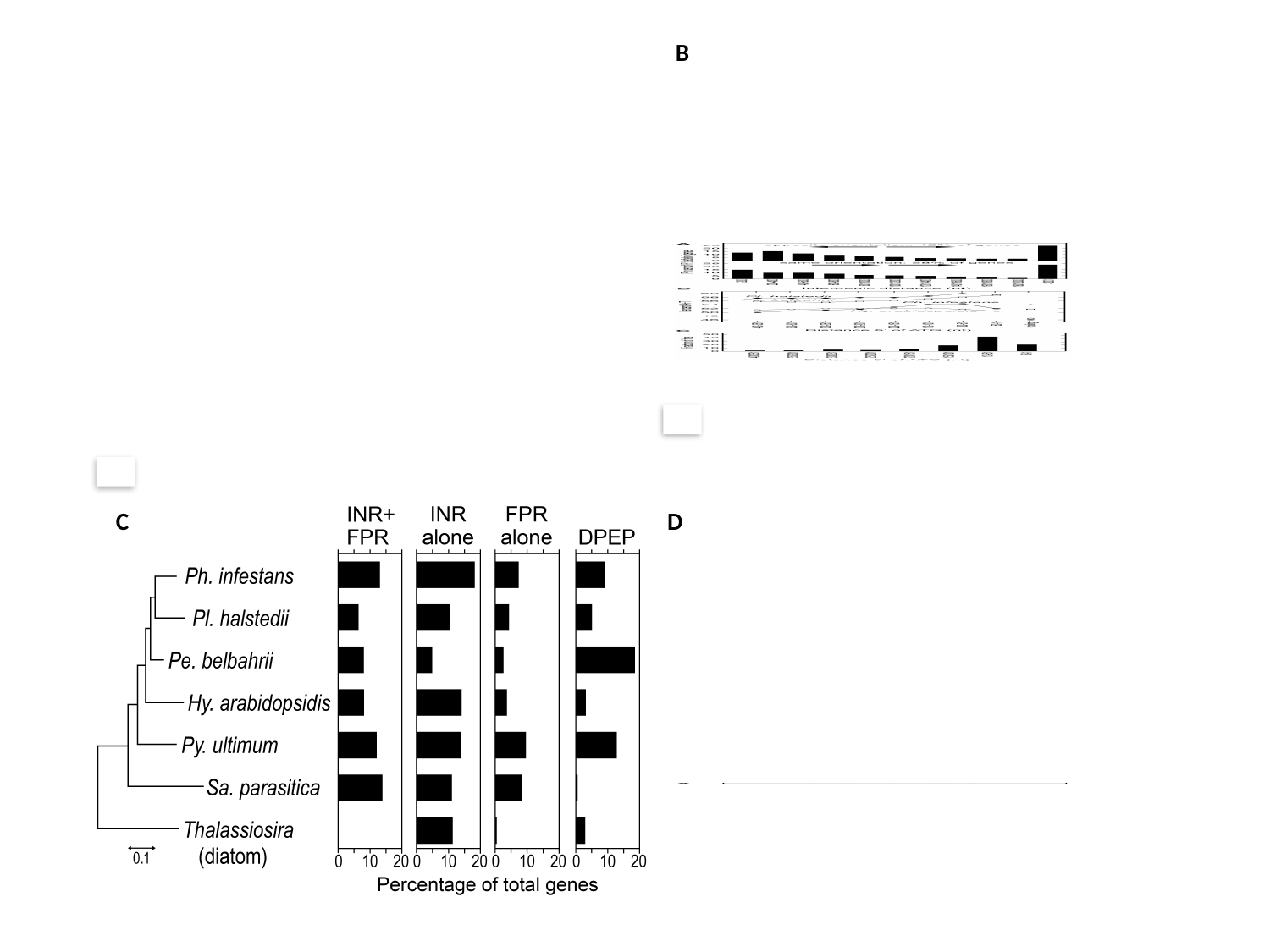

B
B
B
C
D

### Supplementary_Figure_4_photo_lyases.pptx

#### Slide 1
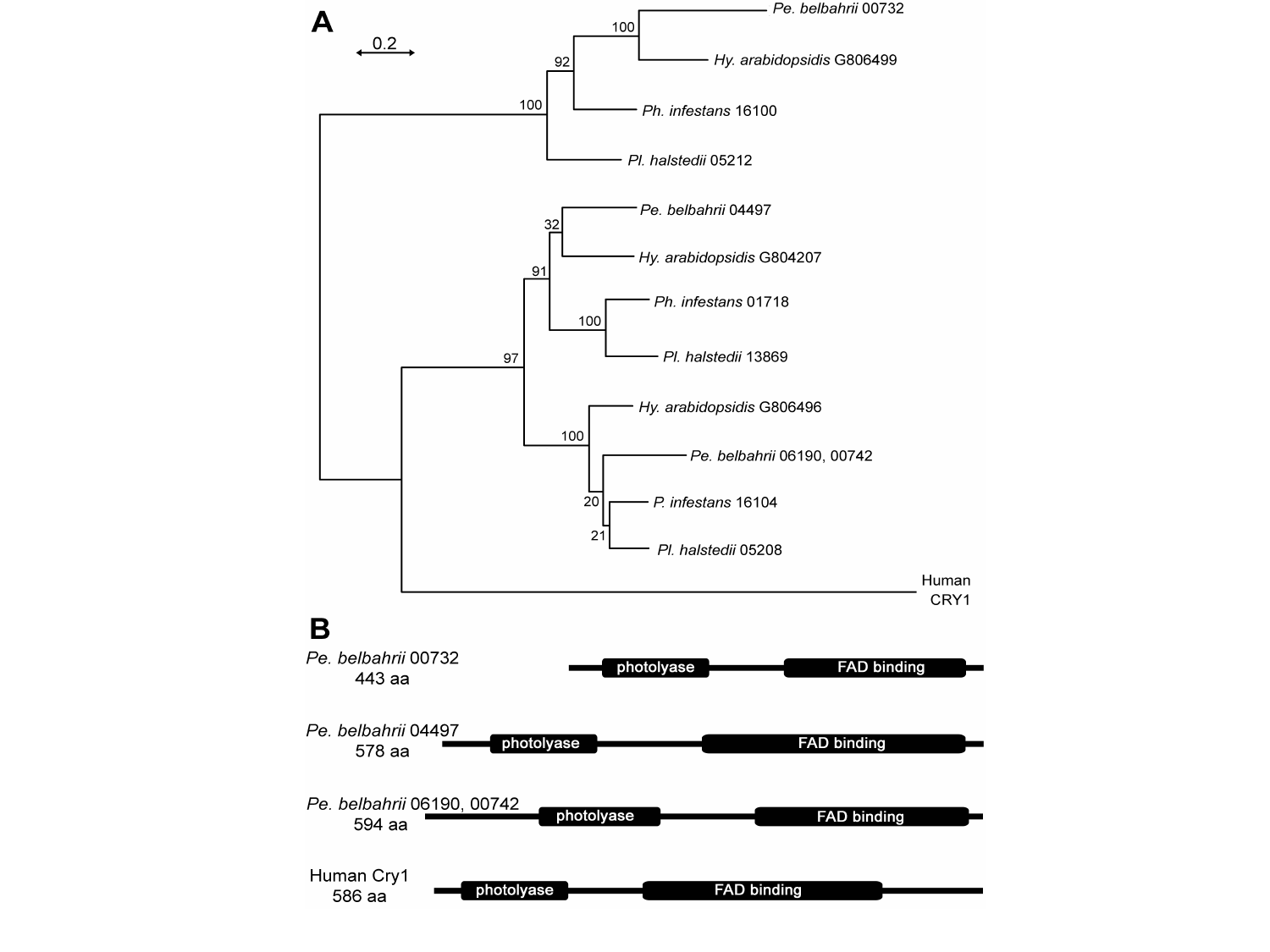
